# Wheat avenin-like protein and its significant Fusarium Head Blight resistant functions

**DOI:** 10.1101/406694

**Authors:** Yujuan Zhang, Xinyou Cao, Angela Juhasz, Shahidul Islam, Pengfei Qi, Maoyun She, Zhanwang Zhu, Xin Hu, Zitong Yu, Steve Wylie, Mirza Dowla, Xueyan Chen, Rongchang Yang, Xianchun Xia, Jingjuan Zhang, Yun Zhao, Nan Shi, Bernard Dell, Zhonghu He, Wujun Ma

## Abstract

Wheat Avenin-like proteins (TaALP) are atypical storage proteins belonging to the Prolamin superfamily. Previous studies on ALPs have focused on the proteins’ positive effects on dough strength, whilst no correlation has been made between TaALPs and the plant immune system. Here, we performed genome-wide characterization of ALP encoding genes in bread wheat. In silico analyses indicated the presence of critical peptides in TaALPs that are active in the plant immune system. Pathogenesis-related nucleotide motifs were also identified in the putative promoter regions of TaALP encoding genes. RT-PCR was performed on *TaALP* and previously characterised pathogenesis resistance genes in developing wheat caryopses under control and *Fusarium graminearum* infection conditions. The results showed that *TaALP* and NMT genes were upregulated upon *F. graminearum* inoculation. mRNA insitu hybridization showed that *TaALP* genes were expressed in the embryo, aleurone and sub-aleurone layer cells. Seven *TaALP* genes were cloned for the expression of recombinant proteins in *Escherichia coli*, which displayed significant inhibitory function on *F. graminearum* under anti-fungal tests. In addition, FHB index association analyses showed that allelic variations of two ALP genes on chromosome 7A were significantly correlated with FHB symptoms. Over-expression of an ALP gene on chromosome 7A showed an enhanced resistance to FHB. Yeast two Hybridization results revealed that ALPs have potential proteases inhibiting effect on metacaspases and beta-glucosidases. A vital infection process related pathogen protein, *F. graminearum* Beta-glucosidase was found to interact with ALPs. Our study is the first to report a class of wheat storage protein or gluten protein with biochemical functions. Due to its abundance in the grain and the important multi-functions, the results obtained in the current study are expected to have a significant impact on wheat research and industry.

## Introduction

Plants have evolved an immune system to recognize and respond to pathogen attack (*1*). Initially, transmembrane receptors on the cell surface detect and recognize the pathogen via pathogen-associated molecular patterns (PAMPs). Adapted pathogens can suppress the PAMP-triggered immunity (PTI) by releasing effector molecules into host plant cells. Plants, in turn, activate a second line of defence, the effector-triggered immunity (ETI) that represses action of the effector molecules (*1*). Pathogen-infected tissues generate a mobile immune signal consisting of multiple proteins as well as lipid-derived and hormone-like molecules, which are transported to systemic tissues, where they induce systemic acquired resistance (SAR) (*2*). SAR is associated with the systemic reprogramming of thousands of genes to prioritize immune responses over routine cellular requirements (*3*). Diverse hormones, such as salicylic acid (SA), jasmonic acid (JA), ethylene (ET), and abscisic acid (ABA) as well as other small phytohormones, play pivotal roles in regulation of this defence network (*4-7*). The signalling pathways cross-communicate in an antagonistic or synergistic manner, providing the plant with a powerful capacity to finely regulate its immune response (*5, 6*). Resistance in plants against pathogen attack can be acquired by resistance genes that biosynthesize metabolites and proteins that directly suppress and/or contain the pathogen to initial infection through their antimicrobial and/or cell wall reinforcement properties. Resistance is achieved specifically by the recognition of pathogen elicitors with plant host receptors, resulting in the induction of signalling events that include changes in ion fluxes, phosphorylation and production of proteins and reactive oxygen species (*8, 9*).

Bread wheat (*Triticum aestivum)* is the third most cultivated crop worldwide, and a major source of daily calories for the human population (*10*). *Fusarium graminearum* is a “hemibiotrophic” pathogen capable of causing wheat head and seedling blight, resulting in yield loss and trichothecene mycotoxin contamination, which is toxic to humans and animals (*11, 12*). In many Asia countries including China, the FHB is referred as wheat cancer in recent years. Several historical wheat growing zones have ceased wheat production due to severe FHB disease. The disease is now fast expanding to wheat growing zones that no FHB disease was occurred in the past. Breeding wheat varieties resistant to FHB has become one of the most important tasks. Better knowledge of the defence mechanisms and genetic engineering provides an effective approach to improve wheat resistance to the disease during breeding. Proteomics approaches have revealed that *F. graminearum* produces extracellular enzymes, such as lipases, xylanases, pectinases, cellulases and proteases (*13-15*), and other proteins, such as hydrophobins, small cysteine rich proteins. These proteins or enzymes may act as pathogenicity factors in plant–microbe interactions (*13*). Proteome studies on *F. graminearum* infected wheat spikes revealed that proteins could be involved in antioxidant, JA, and ethylene (C_2_H_2_-type) signalling pathways, phenylpropanoid biosynthesis, antimicrobial compound synthesis, detoxification, cell wall fortification, defence-related responses, amino acid synthesis, and nitrogen metabolism (*16, 17*). While various transcriptome studies have identified differentially expressed genes of resistant and susceptible wheat spikes infected with *F. graminearum*, suggesting that FHB resistance is conferred by multiple genes (*18-21*). Further, the defence related genes were functionally catalogued to different classes based on previous patho-transcriptomic studies, such as transcription and signalling related genes and hormone (auxins, gibberellins, ABA and SA) metabolism related genes (*20, 22-25*); cysteine-rich antimicrobial peptides (AMPs) (*26-28*); GDSL-lipases (*29*); proteolysis including serine proteases (*30*); peroxidases (POD) (*31*); genes related to cell wall defence (*32-35*), secondary metabolism and detoxification involved genes; Toll-IL-IR homology region (*36*), and miscellaneous defence-related genes, ie., disease resistance-responsive family protein (*37*), NBSLRR disease resistance protein (*38*).

Protein classification according to their conserved domains give insights into sequence and structural and functional correlations. According to the Pfam analysis, many wheat grain specific proteins belong to the prolamin superfamily (http://pfam.xfam.org/). Among them, proteins with LTP-2 (*39, 40*), Tryp-alpha-amyl domain (*41, 42*) and Hydrophobic-seed domain (*43*) were reported to be involved in the plant immunity system and have protease inhibition and antifungal activities. Proteins with a gliadin domain, including the gamma gliadin, LMW glutenin, alpha gliadin, puroindoline, and avenin-like protein (ALP), have been considered as typical storage proteins and have not previously identified with biochemical functions. Their known biological role is as nutrient reservoirs for seed germination. As most storage proteins, the ALPs also have positive effects on wheat flour and dough quality (*44-47*). The current study reports for the first time the molecular characterisation and functional study of *TaALP* in the aspects of anti-fungal activities. Results clearly demonstrated that the ALPs belong to a pathogen-induced prolamin superfamily member gene family. It possesses significant function in resistant to the infection of the FHB pathogen *F. graminearum*. It is expected that the ALPs’ FHB resistant function can be efficiently utilised in controlling FHB. Identifying the potential linkage between *ALPs* and the underlying mechanisms of a range of the newly identified FHB resistant gene and QTLs may further enable successful control of FHB.

## Materials and methods

### Plant Materials

A natural population comprised of 240 wheat cv. s or accessions was used to evaluate the allelic effects on FHB resistance. Eleven lines were sourced from CIMMYT (Centro Internacional de Mejoramiento de Maíz y Trigo). The other 229 lines were from different provinces of China. A double haploid (DH) population Yangmai-16 x Zhongmai-895 consisting of 198 lines were also used for field inoculation assays. Australia premium bread wheat cultivar Mace and Spitfire, and Mace×Spitfire DH line 241 were used for a glasshouse inoculation study in Murdoch University.

### Instruments and Reagents

A HPLC analysis was conducted with an Agilent Series 1200 liquid chromatograph equipped with a quaternary gradient pump system and a diode array UV-Vis detector system connected to a reversed-phase (RP) SB-C 18 column (5 μm, 4.6 × 250 mm, Agilent, USA). Data collection was performed using ChemStation software (Agilent, USA). Chromatograph-grade acetonitrile were purchased from Sigma-Aldrich Co. Ltd (St. Louis, Missouri, USA). Dithiothreitol (DTT), trifluoracetic acid, Sinapinic acid (SA), NaI, ammonium acetate (NH_4_Ac), methanol (MeOH), 4-vinylpyridine (4VP), guanidine HCl, TRIS were purchased from Sigma-Aldrich Co. Ltd. Reagents were of analytical grade and dissolved in deionized water (18 MΩ cm). acrylamide stock solution (30% acrylamide: 0.8% bis acrylamide; Cat #161-0154, Bio-Rad Laboratories, Hercules, CA, USA) with 4.2 mL of water, 3 mL of 3 M Tris-HCl (pH 8.8), 120 μL of 10% SDS, 120 μL of 10% ammonium persulfate (APS) and 6 μL of tetramethylethylenediamine (TEMED).

### Preparation of Albumin and Globulin Protein extracts

Australia spring bread wheat varieties Mace and Spitfire were used for albumin and globulin protein extraction. The albumin/globulin proteins were extracted from 100 mg of flour according to the procedure of Dupont *et al*. (*48*). Briefly, 100 mg of flour was extracted with 1 mL of 0.3 M NaI, 7.5% 1-propanol (NaI-propanol), and centrifuged at 4500 g for 10 min, After two extractions, the supernatant fractions were pooled in 15 ml tubes, precipitated with four volumes of ice-cold (-20°C) NH_4_Ac-MeOH (0.1 M ammonium acetate in 100% methanol), stored at -20 °C for at least 48 h, and centrifuged as above. The supernatant fluid was transferred into 50 ml tubes and precipitated with four volumes of ice-cold acetone and incubated at -20 °C overnight. Following incubation, the fluid was centrifuged as above to yield albumin/globulin fraction pellets.

### Determination of Individual ALPs of wheat cv. Mace and Spitfire via RP-HPLC Analysis coupled with SDS-PAGE, MALDI-TOF, and LC/MS

#### RP-HPLC

Freeze-dried protein pellets were dissolved in 500 µL 6 M guanidine HCl (with a concentration of 1 mg mL^-1^) adjusted to pH 8.0 with TRIS, plus 50 mM DTT, and then alkylated with 4-vinylpyridine (4VP), prior to HPLC analysis (*49*). Albumin and globulin proteins extracted from Spitfire and Mace seeds were analyzed by RP-HPLC. A linear elution gradient was performed using two mobile solvents: the polar solvent A consisting of 0.1% trifluoroacetic acid (TFA), (v/v) in type I ultrapure water (18 MΩ·cm specific resistance) and the non-polar solvent B consisting of 0.1% TFA (v/v) in acetonitrile (ACN). Absorbance was monitored at a detection wavelength of 210 nm, and the flow rate was kept at 0.6 mL min^−1^. The elution gradient conditions were set as follows: from 0 to 51 min, eluent B was increased from 20% to 60%; from 51 to 53 min, eluent B was increased from 60% to 80% and then maintained at 80% for 5 min for washing the column, then decreased to the starting B concentration in 1 min and maintained for 10 min for the next run. The injection volume was 100 µL. The proteins eluted from individual peaks were collected with reference to the chromatographic profile captured in real time and pooled from three runs. RP-HPLC chromatographic finger print profiles showed no variation between runs, thus the elution of each run could be combined to increase the amount of protein in the final sample for later analysis. Samples were immediately frozen at −80 °C for 24 h and lyophilized. Lyophilized samples were stored at room temperature before MALDI-TOF and SDS-PAGE analyses.

#### SDS-PAGE

To identify the ALPs from RP-HPLC eluates, SDS-PAGE was used to separate the protein mixtures of each RP-HPLC eluate, and SDS-PAGE bands of interest were cut for protein peptides sequencing. Then, 12% SDS-PAGE was prepared following Fling and Gregerson’s method (*50*). Briefly, the gel comprises two layers: the separating layer and the stacking layer. The separating gel was prepared by mixed 4.2 mL of acrylamide stock solution (30% acrylamide: 0.8% bis acrylamide; Cat #161-0154, Bio-Rad Laboratories, Hercules, CA, USA) with 4.2 mL of water, 3 mL of 3 M Tris-HCl (pH 8.8), 120 μL of 10% SDS, 120 μL of 10% ammonium persulfate (APS) and 6 μL of tetramethylethylenediamine (TEMED). After polymerization, the separating gel was layered with the stacking gel prepared using 1 mL of acrylamide solution, 750 μL 1 M Tris-HCl (pH 6.8), 4.25 mL of water, 60 μL of 10% SDS, 60 μL 10% APS, and 4 μL of TEMED. Pelleted samples of HPLC eluates described above were mixed with 10 μL 2×laemmli sample buffer SDS loading buffer (Bio Rad).

Electrophoresis was carried out in a modified Laemmli system (*51*). Runs were performed with running buffer of 25 mM Tris-HCL, 192 mM glycine and 0.1% SDS at 120 volts for 2 h. The gels were stained in Coomassie Brilliant Blue (CBB) solution (R-250). Protein standards (Bio-Rad) were used to estimate the molecular size of the proteins. The gels were scanned by a gel Proteomic Imaging System, “Image lab 5.0” (Bio-Rad).

#### MALDI-TOF

MALDI-TOF-MS was used to obtain the mass spectra profile of albumin/globulin fractions obtained from individual HPLC peaks (fractions) with and without 4VP alkylation. The albumin/globulin fraction protein extracts were prepared for MALDI-TOF-MS test, whereas the pelleted RP-HPLC eluted protein samples were diluted 20 times for MALDI-TOF-MS test. Each individual RP-HPLC eluates were lyophilized, the freeze-dried eluates were dissolved with 10 µL ultrapure water, 1 µL was used for MALDI-TOF-MS, and the residues were saved for SDS-PAGE running. Sample preparation was carried out according to the dried droplet method (*52*), using sinapinic acid (SA) as matrix. The matrix solution was prepared by dissolving SA in ACN/H_2_O/MeOH (60:8:32 v/v) at a concentration of 20 mg mL^-1^. All samples, including the RP-HPLC eluates, the raw albumin/globulin extracts and the alkylated albumin/globulins extracts were mixed with SA at the ratio of 1:9 (v/v) individually, and firstly, 1 µL of this protein-SA mixture was deposited onto a 100-sample MALDI probe tip. After drying, another 1 µL of this protein-SA mixture was added, then dried at room temperature. The mass spectra for each sample was recorded on a Voyager DE-PRO TOF mass spectrometer (Applied Biosystems, Foster City, CA, USA) using a positive linear ion mode at an accelerating voltage of 25 kV and a delay time of 700 ns by capturing 1000 spectra of a single laser shot with a mass range of 15000-45000 m/z.

#### Protein identification by MS/MS

Protein bands of interest were manually excised from gels and analysed further by mass spectrometric peptide sequencing. The spots were analysed by Proteomics International Ltd. Pty, Perth, Australia. Protein samples were trypsin digested and the resulting peptides were extracted according to standard techniques (*53*). Tryptic peptides were loaded onto a C18 PepMap100, 3 μl (LC Packings) and separated with a linear gradient of water/acetonitrile/0.1% formic acid (v/v), using an Ultimate 3000 nano HPLC system. The HPLC system was coupled to a 4000Q TRAP mass spectrometer (Applied Biosystems). Spectra were analysed to identify the proteins of interest using Mascot sequence matching software (Matrix Science) with taxonomy set to *Viridiplantae* (Green Plants). All searches used the Ludwig NR. The software was set to allow 1 missed cleavage, a mass tolerance of ± 0.2 Da for peptides and ± 0.2 for fragment ions. The peptide charges were set at 2+, 3+ and 4+, and the significance threshold at P < 0.05. Generally, a match was accepted where two or more peptides from the same protein were present in a protein entry in the *Viridiplantae* database.

### Derivation of Coding Sequences Obtained from *T. aestivum*

Cleavages of signal peptides were predicted using the SignalP 3.0 Server (http://www.cbs.dtu.dk/services/SignalP/). MW’s and pI’s of deduced proteins were calculated using the Protein Parameter tool (http://web.expasy.org/protparam/) found on the ExPASy Proteomics Server. Sequence alignments were performed using ClustalW2 (http://www.ebi.ac.uk/Tools/msa/clustalw2/) with default settings.

### Disease screening

A combination of Type I and Type II FHB resistances was assessed in field nurseries at the Nanhu Experiment Station, Food Crops Institute, Hubei Academy of Agricultural Sciences, (Wuhan, Hubei Province, China) during 2013-2014, 2015-2016, and 2016-2017 crop seasons. The materials used for the 2013-2014 and 2015-2016 crop seasons were 240 wheat lines collected nation-wide in China, while the materials used for the 2016-2017 crop season were the 198 DH lines of Zhongmai 985 x Yangmai 16. The experiments were carried out in randomized complete block designs with two replications. Each plot comprised double 1 m rows with 25 cm between rows. An overhead misting system was applied to favour *Fusarium* infection and development. Plots were spray-inoculated at a concentration of 50,000 spores/ml at anthesis, when 50% of the spikes in the plot were flowering. Conidial inoculum comprised a mix of two highly aggressive isolates of *F. graminearum* isolated from Huanggang and Wuhan, Hubei Province. Ten spikes from different plants in each plot were labeled with blue tape to facilitate scoring. These spikes were assessed 21 DAP for incidence (percentage of diseased spikes) and severity (percentage of diseased spikelet on infected spikes). The FHB index was calculated using the formula FHB index (%) = (Severity × Incidence)/100 (*54*). Naturally occurring FHB was assessed during the 2016-2017 crop season, and the plots were assessed 20, 24, and 28 DAP based on evaluation of FHB index of the plots.

### Promoter analysis

Biotic defense related transcription factor binding sites (ATCAT, TGACG, TTGAC, CANNTG) and TF specific binding sites related to biotic defense were collected from public promoter motif and TF databases (Plant TFDB - http://planttfdb.cbi.pku.edu.cn; plantCARE - (http://bioinformatics.psb.ugent.be/webtools/plantcare/html/) and PLACE - https://sogo.dna.affrc.go.jp) and used for transcription factor binding site prediction on 1100 bp nucleotide sequences including 1000 bp promoter region upstream from the start codon of the avenin-like protein coding genes. Promoter sequences were retrieved from the *Triticum aestivum* cv. Chinese Spring (CS42) whole genome assembly (Triticum_aestivum_CS42_TGAC_v1, Earlham Institute, UK). The TF binding motifs were annotated according to their hormone and TF family specificity. Promoter motifs were mapped using the CLC Genomics Workbench v. 11 (CLCBIO Aarhus, Denmark) both onto sense and anti-sense strands with 100% sequence identity. TFBSs belonging to the same annotation group were marked with the same colour.

### Point inoculation on wheat spikelets

Glasshouse based experiments were carried out at Murdoch University, glasshouse 2. The *F. graminearum* strain was sourced from Curtin University. The *F. graminearum* isolates were grown on mung bean agar plates (MBA) for four weeks to produce spores. Spores were collected via flooding of the cultures with sterile water, and the spore concentration in the suspension was adjusted to 5×10^5^ conidia/mL before point inoculation. Point inoculation of wheat spikelets was performed as follows: inoculation of 10 μL spore suspension/deionized water into the two-central opposite wheat flowering spikelets, which were then covered in a polythene bag for 48-72h to maintain a high humidity. Infected spikelets were counted after two weeks. Mock inoculation was done by replacing spore solutions with sterile deionised water and treating spikelets in the same way. Inoculation experiments were repeated three times independently. Infected and mock spike samples were collected 7, 13, and 42 DAP. After sampling, plant material was immediately frozen in liquid nitrogen and stored at -80°C until use. For each biological replicate, two inoculated spikes per time point were collected and for each biological replicate, three technical replicates were conducted.

### Overexpression of TaALP-bx-7AS gene in transgenic wheat lines

An agrobacterium mediated gene transformation procedure was followed to overexpress a *TaALP* gene on chromosome 7A (*TaALP-bx-7AS*) in cv. Fielder. The T2 plants were screened for FHB symptoms under combined Type I and Type II FHB inoculation in glasshouse. The symptom scoring procedure was the same as that used in the field nursery.

### Characterization of genes in Spitfire and Mace and marker design

Genomic DNA of wheat cv. s was extracted from 1-week-old seedlings using the cethyltrimethyl ammonium bromide (CTAB) method as reported (*55*). Based on the *TaALP* gene sequence from Juhasz et al (2018), specific primers were designed for different loci. These amplified the full gene from the 5’ and 3’ ends. The PCR products were separated on 1.5% (w/v) agarose gels, and e bands of the expected size purified using a Gel Extraction Kit (Promega). Subsequently, the purified PCR products were amplified using BigDye@version 3.1 terminator mix (Applied Biosystems) and sequenced. RefSeq v1.0 gene models in Chinese Spring were used to analyse sequences. Further specific primers were designed for each hit chromosome using Primer V5.0 software (http://www.premierbiosoft.com) (Chapter 2).

### RNA isolation

Total RNA was extracted using TRIzol reagent (Invitrogen Canada, Inc., Burlington, Ont., Canada, catalogue No. 15596026) according to the manufacturer’s protocols. cDNA was synthesized using an RNA reverse transcription kit (Bioline, London, UK, Catalogue No. BIO-65053). qRT-PCR was performed on a Rotor-Gene RG3000A detection system (Corbett Research) using SensiFAST SYBR No-ROX Kit (Bioline, London, UK, Catalogue No. BIO-98005) as follows: hold at 95°C for 2 min, followed by 45 cycles of 95°C for 10s, 60°C for 15s, 72°C for 30s. A melting curve was performed to determine the specificity of each PCR primer by incubating the reaction at 95°C for 20 s, cooling at 55°C for 10 s, and increasing to 95°C at a rate of 0.5°C/10 s. The reference gene *β*-actin was used for the normalization of all qRT-PCR data. The 2^-ΔΔCt^ method (*56*) was used to calculate the relative expression levels with three technical repeats (Chapter 2).

### *In situ* hybridization

To generate gene-specific anti-sense probes, a 750-bp and a 500-bp *TaALP* cDNA clone, pspt19 (RGRC-NIAS; http://www.rgrc.dna.affrc.go.jp/stock.html), was digested with *Bam*HI and *Sac*I, respectively, and transcribed in vitro under the T7 and SP6 promoters with RNA polymerase using the DIG RNA labeling kit (Sigma Aldrich). *In situ* hybridization was performed according to the protocol of Kouchi and Hata (*57*) (**Appendix Table 3**).

### Recombinant TaALP production

Full-length *TaALP* cDNA was inserted into the bacterial expression vector pET28a (+) (Novagen), and the constructs were then introduced into *Escherichia coli* BL21(DE3) codon plus. Bacteria contain the plasmids were grown in Luria-Bertani (LB) medium containing 50μg/ml kanamycin at 37°C to OD_600_=0.6. Expression of the fusion protein His-ALP was induced by addition of 1 mM isopropyl *β*-D-1-thiogalactopyranoside (IPTG) and incubation at 25°C for 16 hr. Bound proteins were eluted with sodium phosphate buffer containing increasing concentrations of imidazole and detected by 12% SDS-PAGE. Nonspecific proteins purified from the bacteria with the pET28a (+) vector were used as control (**Appendix Table 3**).

### *In vitro* antifungal activity of recombinant ALPs

An agar-gel diffusion inhibition assay was carried out in order to determine the *in vitro* anti-fungal activity for inhibition of mycelial growth of *F*. *graminearum*. Three 5-mm diameter mycelial disks (3-day-old culture) of the strain was placed in the PDA plate with 100 μl of the recombinant protein sample and incubated at 23 °C for 3 days. Inhibitory zones from different recombinant samples were visually compared with those from the control bacterial extracts. Antifungal activity of ALPs proteins against fungi was assayed by micro spectrophotometry of liquid cultures grown in microtitre plates as described previously (*58, 59*). Briefly, in a well of a 96-well microplate, 10 µl of the protein sample (purification buffer as control) was mixed with 90 µl minimal medium (MM) containing fungal spores at a concentration of 1×10^5^ conidia ml^−1^. Growth was recorded after 24 h incubation at 22°C daily. EC_50_ values (the concentration of the antifungal protein required to inhibit 50% of the fungal growth) were calculated from dose–response curves with two-fold dilution steps (*59*). The absorbance was recorded at 595 nm in a 96-well plate reader (Biorad).

### GAL4-based yeast two-hybrid assay

TaALP protein interactions were studied using GAL4-based yeast two-hybrid assay, including protein to protein interactions within the wheat host and these between host and pathogen. A *F*. *graminearum* cDNA library was screened for potential interactions. The *TaALP* gene were amplified using the forward primer *TaALP-NdeI*-F and the reverse primer *TaALP-BamHI*-R (**Appendix Table 3**). The PCR product and plasmid pGBKT7 (CLONTECH Co., United States) were treated with *NdeI* and *BamHI* enzymes (Neb, England), respectively, followed by ligation to construct the recombinant vector pGBKT7:*TaALP-4*. The recombinant vector and the negative control *pGBKT7* were transformed into the wild yeast cells *Y187* (CLONTECH Co., United States), respectively, and cultured in *Trp* lacking media. While prey proteins are expressed as fusions to the Gal4 activation domain (AD) (*60, 61*). The *Ta-MCA* gene and *Ta-NMT* gene were amplified using primers listed in **Appendix Table 3**. The PCR products and plasmid pGADT7 (CLONTECH Co., United States) were treated with *NdeI* and *BamHI* enzymes (Neb, England), respectively, followed by ligation to construct the recombinant vector pGADT7:*TaMCA* and pGADT7:TaNMT. The recombinant vectors were transformed into the wild yeast cells *Y2HGold* (CLONTECH Co., United States), respectively, and cultured in *Leu* lacking medium. The clones grown in the *Leu* lacking medium were mated with the previous *Trp* lacking medium colonies with overnight shaking and then transferred to the *Trp* and *Leu* lacking medium with X-*α*-gal, to allow bait and prey fusion proteins to interact. The DNA-BD and AD are brought into proximity to activate transcription of *MEL1* to test the transcriptional activation activity. Sequences coding for one anti-fungal proteins, *ALP* gene (encoded by *7dyb*, **Appendix >YJ7dyb**), were chemically synthesized according to their amino acid sequences. A metacaspase gene and NMT gene were cloned from a common Australia wheat cv. Lincoln.

### Statistical analysis for the allelic effect

For the allelic effect study, marginal F tests were used to determine the significance of allelic effects on FHB indexes of the 240 wheat varieties (*62*). Markers were nested within the population. The statistical significance of the FHB index was assessed performing T-tests using the SAS/STAT System software, Version 8.0 (SAS Institute Inc. Cary, NG) for the DH population of Yangmai16 x Zhongmai 985. All measurements were carried out in triplicate, and the results presented as mean values ± SD (standard deviation). Statistical analysis was performed via one-way analysis of variance (ANOVA) followed by Duncan’s test. P < 0.05 were considered significant. Data were analyzed using SPSS 19.0 (SPSS Inc., Chicago, IL, USA) for Windows and figures generated using SigmaPlot 12.0 and Photoshop 8.0 for Windows.

## Results

### In silico analyses revealed pathogenesis-related features on ALP encoding genes

To investigate the potential relationship of ALP genes with pathogenesis, the previously characterised pathogenesis-related motifs were retrieved from public database (**Table S1**). A total of 11 motifs, related to different hormones and transcription factor families, were identified. The putative promoter binding regions (1000 bp region upstream the translation starting sites) of 15 *ALP* encoding genes (63) in bread wheat were surveyed for the presence of those motifs (**Figure 1A**). Overall, multiple pathogenesis-related motifs, ranging from 11 to 28, were identified in the promoter binding regions of all *ALP* genes. The 15 wheat *ALP* genes could be divided into 5 orthologous groups: 2 groups of type a (*ax, ay*), 2 groups of type b (*bx, by*), and 1 group of type c (*c*). Interestingly, the highest number (26-28) of pathogenesis-related motifs was observed for *bx* genes, while the lowest was found for *ay* genes, ranging from 11 to 17. When different types of *ALPs* were compared, the highest number of pathogenesis-related motif was observed for type b (146) followed by type a (103), with type c being the lowest (55). In addition, when different chromosomes were compared, the *ALP* genes on 7D have the highest number (108) of pathogenesis related motif, which is higher than these of 7A and 4A (both at 98).

**Figure 1.**
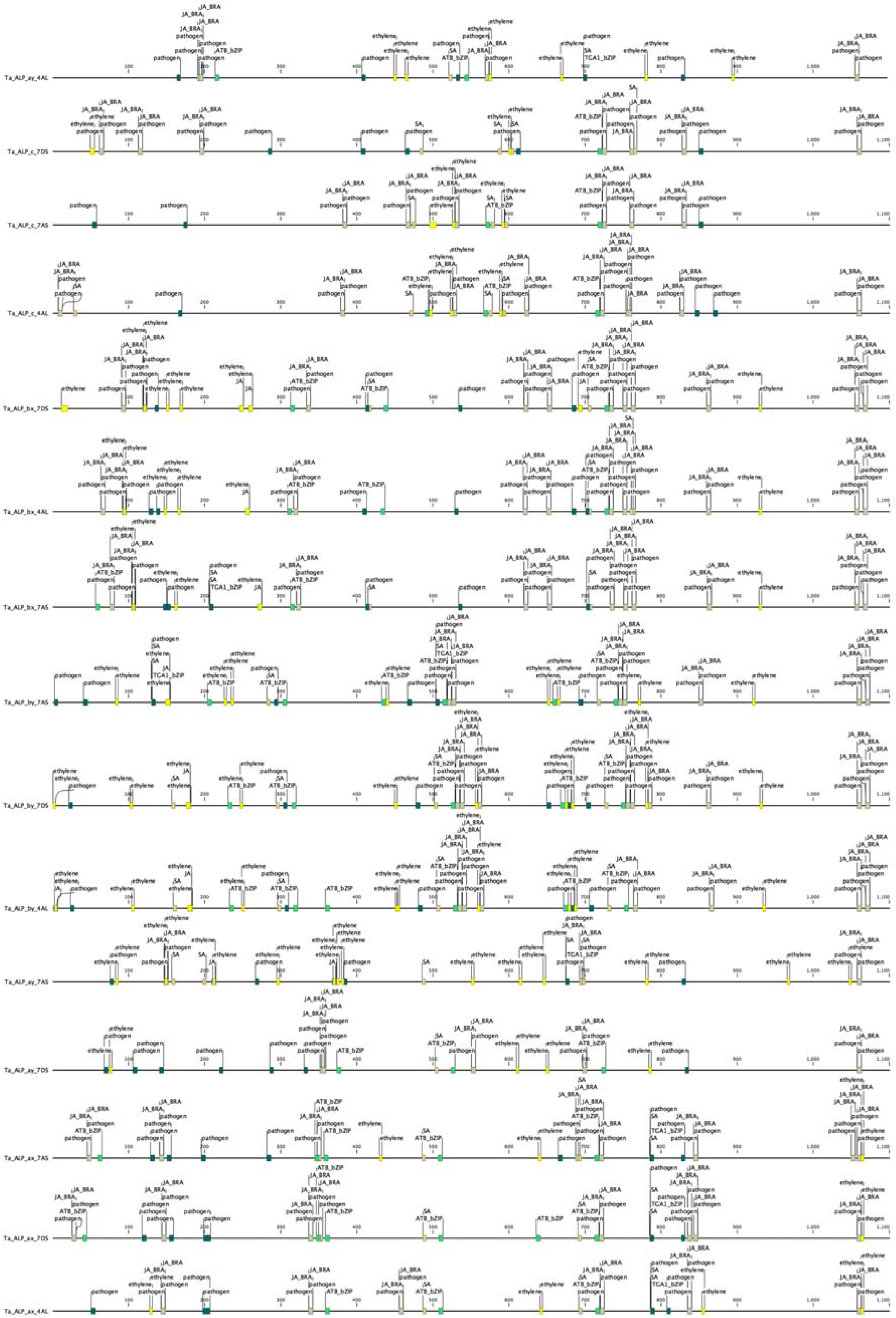
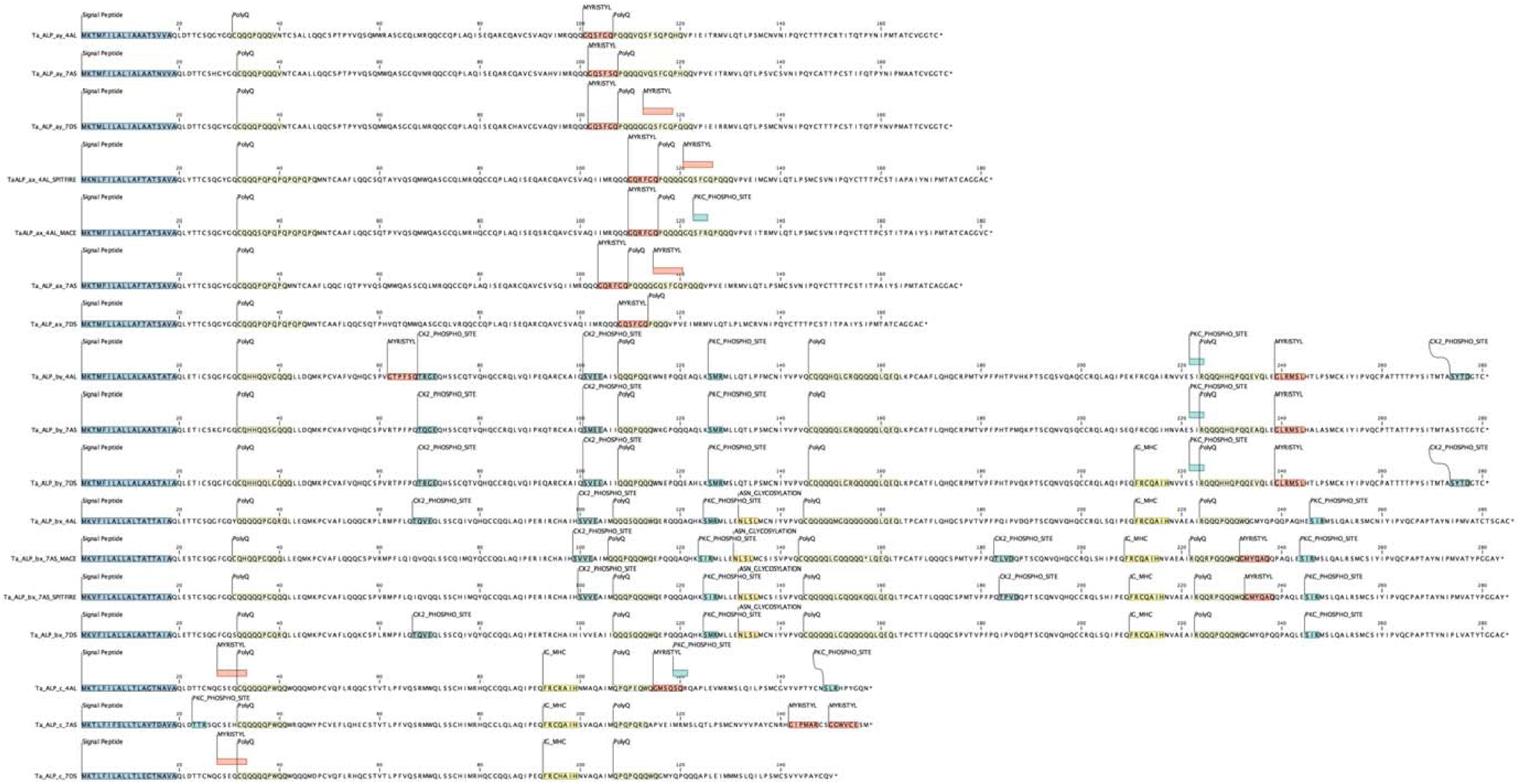
In silico analyses on ALP genes. A. Prediction of pathogenesis-related motif in the promoter regions of ALP genes. B. Prediction of the presence of signal peptides, phosphorylation sites, myristoylation sites, polyQ groups in ALP proteins.

In addition to pathogenesis-related motif analyses in the promoter regions, the predicted amino acid sequences for the 15 *ALP* encoding genes were analysed for the presence of N-myristoylation sites, which have been shown to be related to pathogenesis. In particular, candidate proteins could be cleaved at the myristoylation sites, followed by myristoylation reaction catalysed by N-myristoyltransferse. This process leads to programmed cell death, which confers systemic acquired resistance (SAR). Results showed that 13 out of the 15 ALP proteins contained one or two myristoylation sites (**Figure 1B**), suggesting a potential biological role in pathogenesis resistance.

### Peptide sequencing showed ALPs were cleaved in mature wheat grain

To investigate the content of ALPs in mature wheat grain, total albumin and globulin proteins were extracted from two wheat cultivars, Mace and Spitfire. The presence of ALPs was identified by reverse-phase HPLC (RP-HPLC), SDS-PAGE and Maldi-tof methods. Firstly, the extracted protein samples were separated by RP-HPLC. A total of 36 and 33 elution peaks were identified for Mace and Spitfire, respectively (**Figure S1**). Then, the protein fractions for each HPLC peaks were collected and loaded on SDS-PAGE gel for further separation. As shown in **Figure 2**, most of the collected HPLC fraction contains a mixture of proteins with different molecular weights. The major bands in each fraction were cut out and sent for peptide sequencing. Only those target proteins with molecular size close to or lower than the maximum predicted molecular weight of ALPs (∼ 33 kDa) were analysed. A total of 55 SDS-PAGE bands were sequenced (**Figure 2**; **File S1**). Results (**Figure 2**) showed that 20 and 15 fractions from Mace and Spitfire, respectively, were found to contain ALPs.

**Figure 2.**
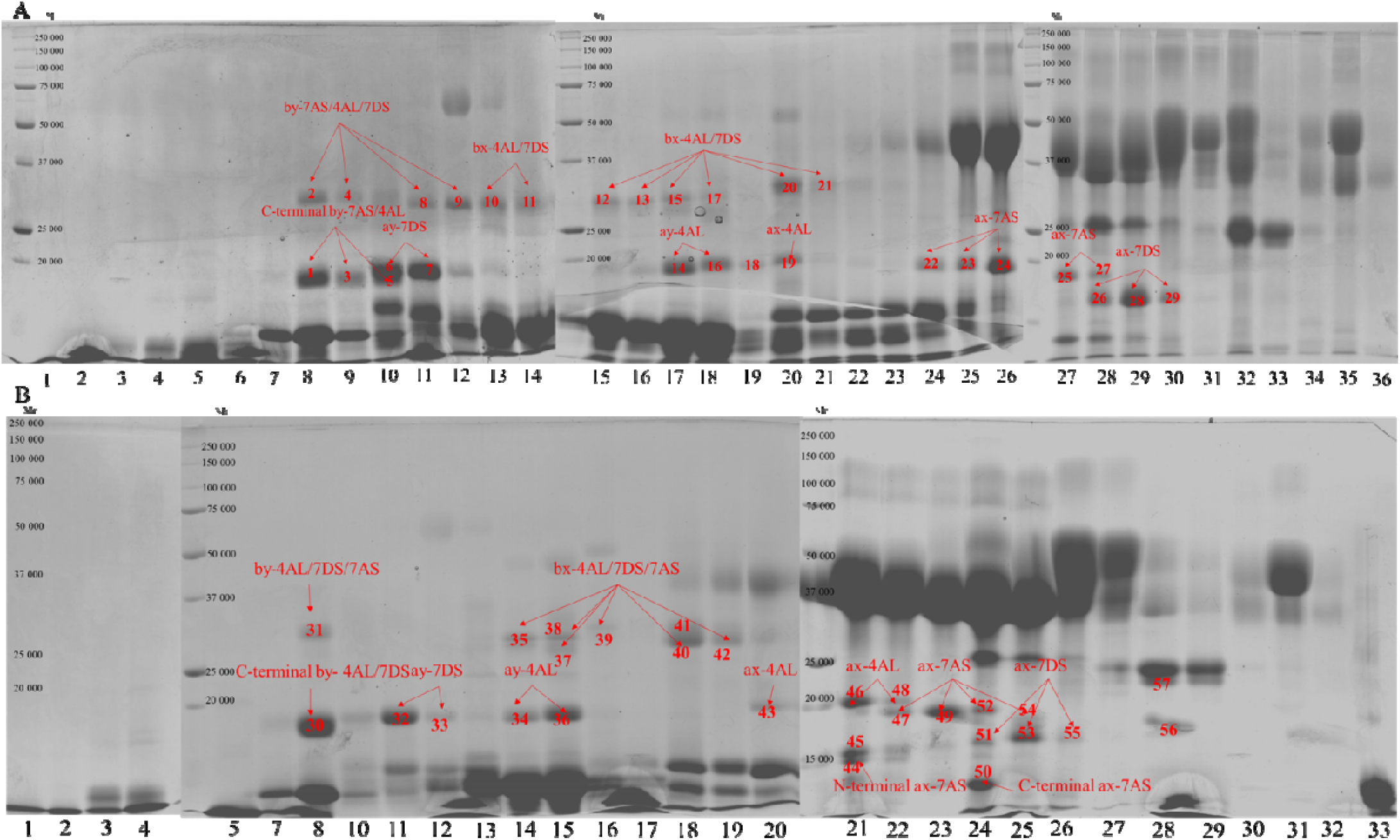
SDS-PAGE gel separation of albumin and globulin proteins.

For Mace, 5 (ay-7DS/4AL, ax-4AL/7AS/7DS) out of the 15 ALPs, belonging to type a, could be identified in fractions 8-11, 17-18, 20, 24-30. These type a ALPs displayed molecular weight similar to the full length ALPs, suggesting an intact form of type a ALPs. In addition, 12 protein bands (2, 4, 8-13, 15, 17, 20-21) were identified as type b ALPs (bx and by) which, however, could not be assigned to specific ALP orthologues. Notably, some identified type “by” ALPs, corresponding to bands 1, 3 and 5, displayed molecular weight at around 18.34 kDa (**Figure S2A**), suggesting an inter-domain cleavage for these ALPs. Other type “by” ALPs, corresponding to bands 2, 4, 8 and 9, contained multiple proteins at ∼ 32.32 kDa and ∼ 28.19 kDa, which were validated by Malti-tof analyses (**Figure S2A, C**). These results suggest the presence of both full length and another kind of partial type “by” ALPs, which may be resulted from the cleavage at the predicted myristoylation sites. This hypothesis is consistent with the predicted molecular weight for myristoylation cleavage and is supported by the peptide sequencing results, which revealed no peptide covering the myristoylation sites. In contrast to the type “by” ALPs, the identified type “bx” ALPs, corresponding to bands 10-13, 15, 17, 20-21, were all characterised as full length ALPs, suggesting that the type “bx” ALPs that do not contain myristoylation sites had no cleavage occurred.

Similar observations were made with Spitfire. Five type a ALPs (ay-7DS/4AL, ax-4AL/7AS/7DS) could be identified in the predicted full length form with molecular weights ranging from 17.90 kDa to 19.20 kDa. Bands 31, 35, 37-42 were identified as type b ALPs, containing both types “by” and “bx”. No type b ALP orthologue could be assigned. For type “by”, molecular weights of 32.43 kDa, 28.28 kDa and 18.41 kDa were observed, suggesting the presence of the intact form and two differently cleaved forms. For type “bx”, molecular weight at 33.01 kDa, 32.92 kDa, 32.67 kDa, 27.61 kDa were identified, indicating the occurrence in the intact and the predicted myristoylation cleaved forms but not in the inter-domain cleavage form. Notable, for both Mace and Spitfire, no type c ALP could be identified in the present study.

### ALP genes were upregulated upon *F. graminearum* inoculation in developing wheat caryopses

To investigate the potential interactions between ALP genes and pathogen resistance, the transcriptional profiles of 7 ALP genes (*ax-7AS/7DS, ay-7DS, by-7AS/7DS, bx-7AS/7DS*), 2 previously characterised anti-virulence gene candidates (*Taxi III, PR.1.1*), and 2 Programmed cell death (PCD) related genes including wheat meta-caspase gene (*TaMCA*4) and N-myristoly Transferase gene (*TaNMT*) were studied by RT-PCR under control and *F. graminearum* inoculation conditions in developing wheat caryopses. A total of 3 wheat lines (Mace, Spitfire, DH line 241) at 3 developmental stages (7 DPA, 13 DPA, 42 DPA) were investigated (**Table 1**). Overall, for the 7 ALP genes, the highest expression was observed at 13 DPA with the exception of 7axb, which was barely expressed at all stages under the control conditions. At 13 DPA, a clear upregulation of ax-7AS, ax-7DS, ay-7DS, by-7AS, bx-7DS and by-7DS upon *F. graminearum* inoculation could be detected in all or some of the three wheat lines. Similar observations could also be made at 42 DPA, when the transcription of ax-7AS, 7axb and 7ayb were significantly upregulated in some wheat lines. Noteworthy, although 7axb is barely expressed in all wheat lines throughout seed development under control condition, significant upregulation of the transcription of this gene was detected at 7 DPA and 42 DPA in DH line 241. In contrast to the *ALP* genes, transcription of *RP.1* and *Taxi III* genes were mainly found at 7 DPA and 42 DPA but not at 13 DPA. At 7 DPA and 42 DPA, clear up-regulation of RP.1 and taxi was observed after *F. graminearum* inoculation, suggesting a positive role for these genes in pathogenesis activities. For MCA and NMT genes, the highest expression occurred at 13 DPA, with very low or no expression at 7 DPA and 42 DPA. At 13 DPA, in contrast to MCA, which displayed variable transcriptional changes among different wheat lines upon *F. graminearum* inoculation, significant up-regulation of NMT was detected in all wheat lines studied.

**Table 1.**
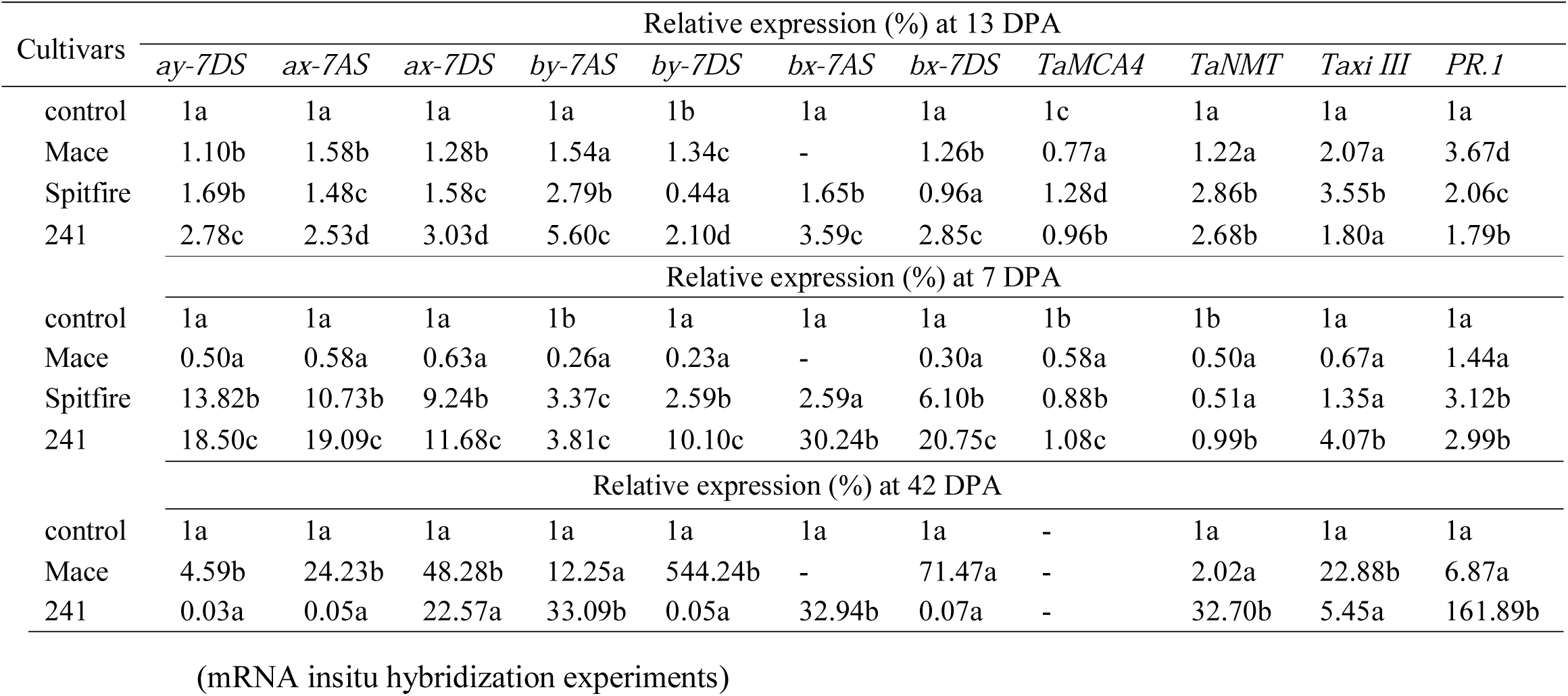
RT-PCR of ALP genes in developing wheat seeds under control and pathogen infection conditions. (mRNA insitu hybridization experiments)

### ALP genes were expressed in the embryo, aleurone, sub-aleurone and transfer cells

To determine the transcriptional domain of ALP genes, mRNA insitu hybridization was performed for two ALP genes: *ay-7DS* and *by-7AS*, representing type a and b, respectively. The developing wheat caryopses of Mace at 15 DPA was used. As shown in Figure 4, clear signals of type a ALP gene *ay-7DS* expression were detected in the embryo (Figure 4A), aleurone cells (Figure 4A), sub-aleurone and transfer cells (Figure 4A). The highest intensity was observed in the aleurone and subaleurone cells, followed by embryo, whilst the signal in the transfer cells is relatively weaker. No signal or very weak signal could be detected in other part of the endosperm, pericarp and husk tissues. Similar results were obtained for type b *ALP* gene *by-7AS*. The transcription of *by-7AS* was observed in embryo (Figure 4B), aleurone (Figure 4B), sub-aleurone (Figure 4B) and transfer cells (Figure 4B), with the highest expression in embryo, aleurone and sub-aleruone cells, whilst relative weaker in transfer cells.

**Figure 4.**
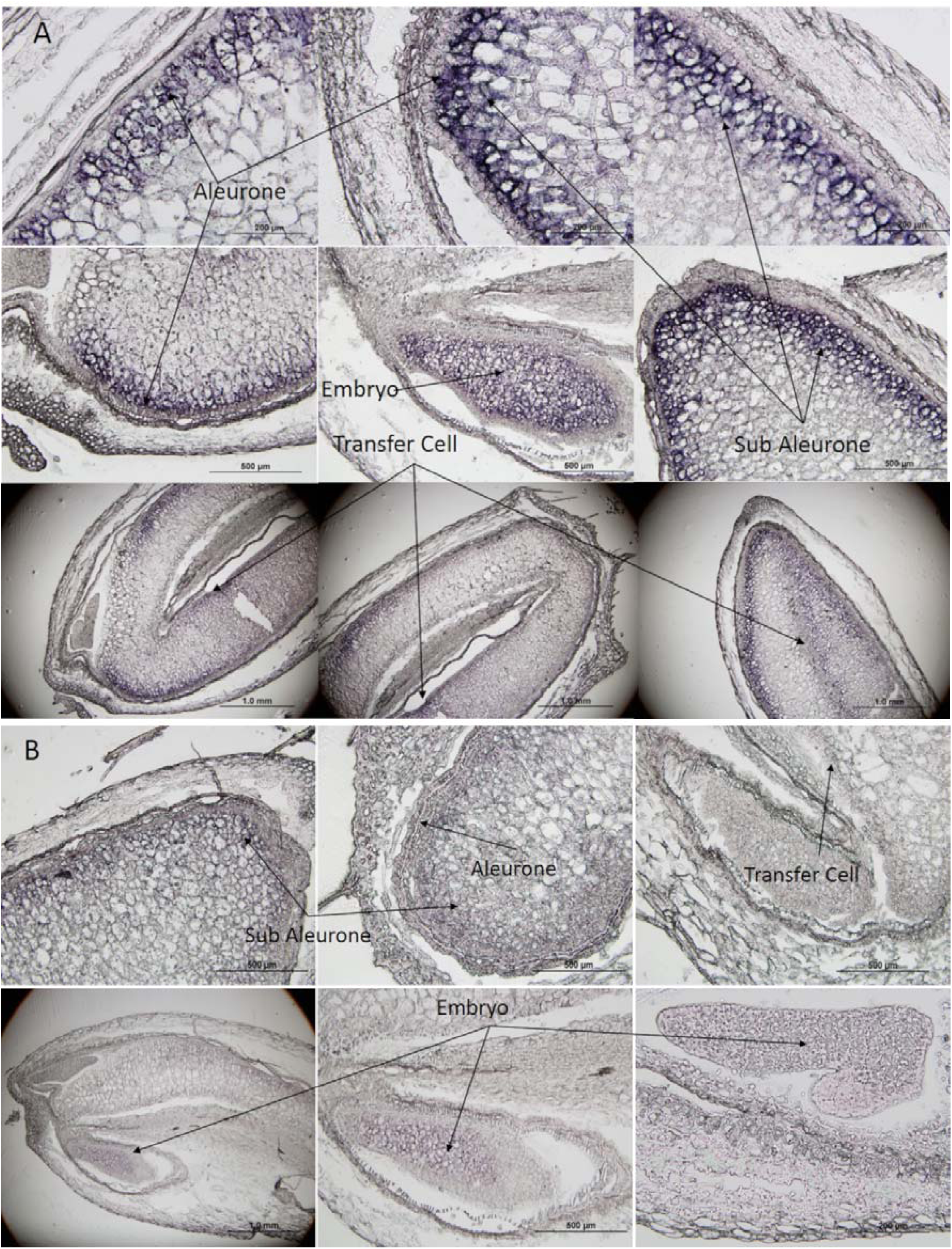
mRNA insitu hybridization on ALP genes in developing wheat caryopses. A. type a ALP; B. type b ALP.

### ALPs displayed significant *in-vitro* anti-fungal function on *F. graminearum*

To determine whether ALP proteins have anti-fungal function, 7 ALP genes (*ay-4AL, ay-7AS*, C-terminal *by-7DS, c-7AS, by-7AS, bx-4AL* and *by-7DS*) were cloned into pET28a(+) vector and induced for recombinant protein production in *E. coli* system. After protein induction, *E. coli* cells were harvested and lysised, followed by centrifugation. The expression of the target ALP proteins in the supernatant solution were confirmed by SDS-PAGE gel (Figure S3). Preliminary anti-fungal tests were performed by studying the inhibitory effects of *F. graminearum* on PDA plates. As shown in Figure S4, compared to the control tests, the growth of *F. graminearum* colonies were clearly inhibited by the recombinant ALP protein solutions, indicating all selected ALPs have anti-*F. graminearum* function. Noteworthy, variable degrees of anti-fungal activity were observed, with ay-4AL, ay-7AS, C-terminal by-7DS displayed the highest and comparable anti-fungal activities, followed by by-7AS. The lowest anti-fungal activities was observed for c-7AS, bx-4AL and by-7DS. Further anti-fungal tests were performed by studying the inhibition effects on *F. graminearum* in minimal medium (MM) media. The E. coli strain harbouring the pET28a(+) vector with no gene insert was used as control. The growth rate of *F. graminearum* was plotted in Figure 5A for the 7 selected ALP proteins. The inhibitory activity of each candidate protein was assessed by calculating the EC_50_ value. Overall, the results are consistent with that obtained from the PDA plate tests. As shown in Figure 5B, ALP, ay-4AL, ay-7AS, and C-terminal by-7DS displayed the lowest EC_50_ values (0.11 – 0.15), suggesting the highest anti-fungal activities for these proteins. This is followed by ALP by-7AS, which has an EC_50_ of 0.42 and demonstrated a moderate anti-fungal activities. The lowest EC_50_ values were observed for ALPs c-7AS, bx-4AL and by-7DS, suggesting these ALPs have relative lower anti-fungal activities toward *F. graminearum*.

**Figure 5A-C.**
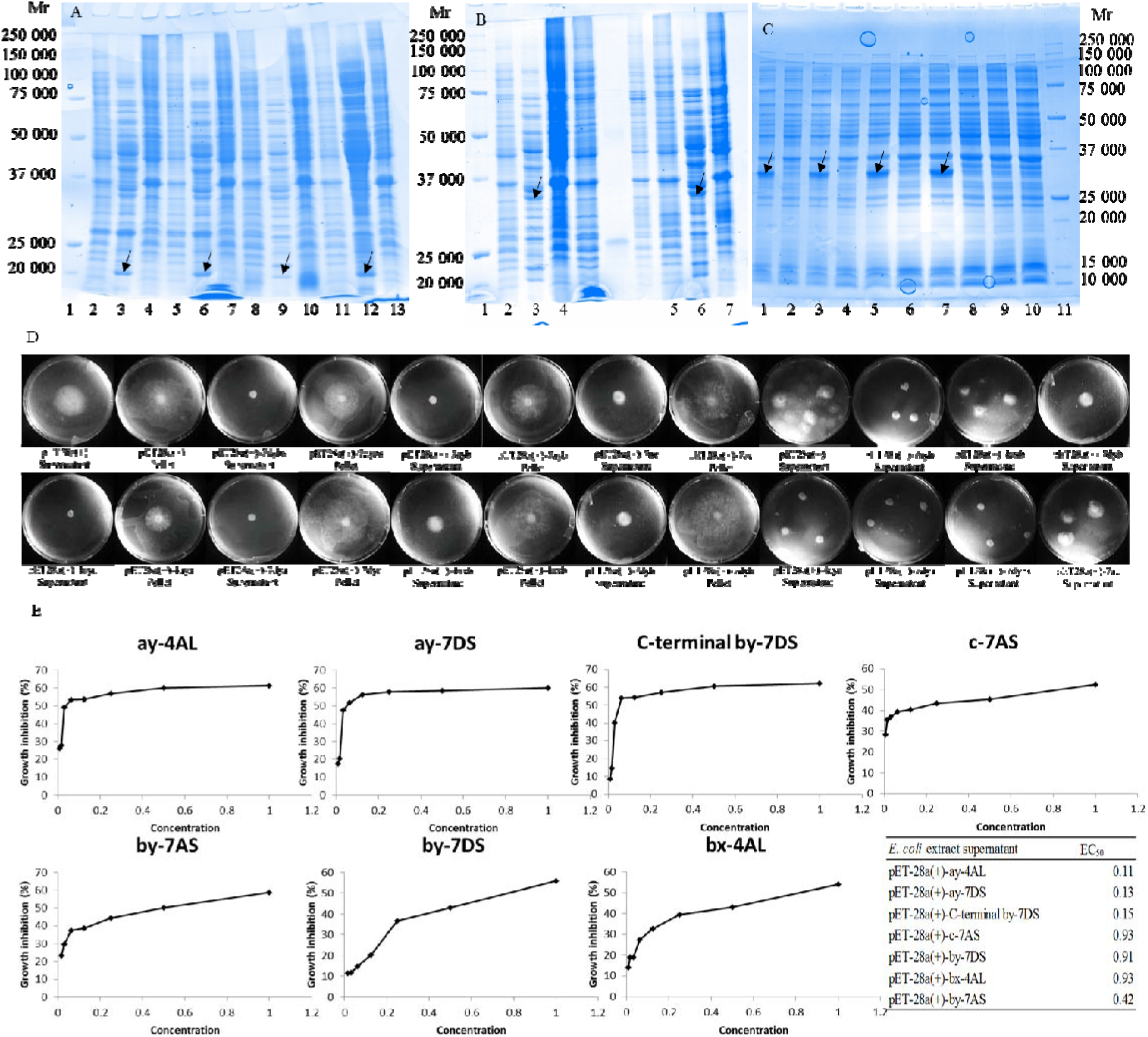
The production of recombinant ALP protein validated by SDS-PAGE gel; D Anti-fungal tests of ALPs on *F. graminearum* on PDA plates; E. Anti-fungal tests of recombinant ALPs on *F. graminearum*. growth rate plot and EC_50_ value calculation.

### ALPs have potential proteases inhibiting effect on metacaspases and beta-glucosidases

In silico, peptide sequencing and gene transcriptional analyses in the present study suggested a positive interaction between ALP genes and pathogenesis-related genes MCA and NMT. To validate the predicted interactions between these proteins, yeast two hybridization experiments were performed. Three ALP proteins (ay-4AL, ay-7AS, C-terminal by-7DS) were selected to study their potential interactions with TaMCA4 and TaNMT proteins. Each gene fragment encoding these corresponding proteins were cloned into both PGADT7 and PGDBKT7 vectors to allow forward and reverse double validations. For each experiment, both vectors, containing one ALP insert and one target pathogenesis-related gene insert, were transformed into yeast strainsY187 and Y2HGold, respectively. The potential protein interactions were assessed by colour reaction on X-alpha-Gal media plate. As shown in Figure 6, all of the three selected ALPs were found to interact with TaMCA4 in both forward and reverse tests. However, those ALPs displayed weaker interactions with TaNMT, which was also confirmed by both forward and reverse interaction tests.

**Figure 6.**
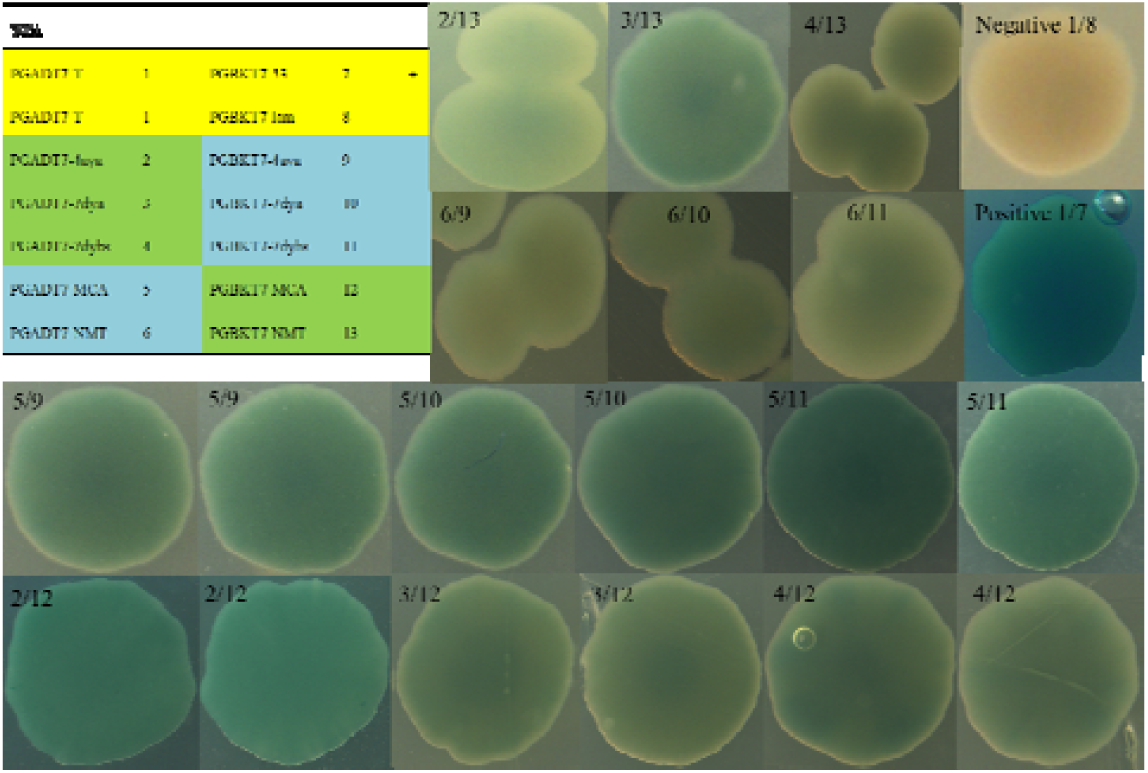
Yeast two hybridization tests of ALPs and pathogenesis-related proteins TaMCA4 and TaNMT.

To further identify the other fungal proteins which may interact with ALP proteins, the above three ALP gene constructs were used to screen the cDNA library constructed from *F. graminearum* strain. Beta-glucosidase which is encoded by a candidate gene (id FG05_01351) was found to interact with all of the three target ALPs. Beta-glucosidase has been shown to be able to hydrolyse the chitin in wheat seed pericarp, which plays an important role during the pathogen infection process. This may explain the molecular basis underlying the anti-fungal function of ALP toward *F. graminearum*.

### Functional ALPs alleles are significantly associated with lower FHB index

To further characterise the potential anti-fungal role of ALPs in wheat, FHB index association analyses were performed to study the ALP allelic effects on 240 wheat cvs. (collected across different regions in China) using one SNP marker (marker ID *bx-7AS*). SNP1 were identified in ALP gene *bx-7AS*, while the other allele resulting in a dysfunctional per-mature termination. The FHB index data were collected from two continuous years for the 240 wheat cvs. grown in two different locations. Results showed that, for the SNP marker, the functional alleles were significantly (*P* < 0.05) associated with a lower FHB index, indicating a positive effects on *F. graminearum* resistance (**Table 2**).

**Table 2.** Statistical analysis of *bx-7AS* gene on FHB index

| Average<br>FHB index | 2013-2014 |  | 2015-2016 |  | 2016-2017 |  |  |  |
| --- | --- | --- | --- | --- | --- | --- | --- | --- |
|  | 21DPA | 21DPA | 21DPA | 21DPA | 21DAP | 20DAP | 24DAP | 28DAP |
| Allele 0 | 77.59 | 80.29 | 73.12 | 75.5 | 57.3 | 29.8 | 44.1 | 55.8 |
| Allele 1 | 71.77 | 74.75 | 63.89 | 65.82 | 50.1 | 22.9 | 34.7 | 45 |
| Increased<br>Rate (%) | -7.5 | -6.91 | -12.62 | -12.83 | -12.57 | -23.15 | -21.32 | -19.35 |
| P value | 0.038 | 0.046 | 0.015 | 0.01 | 0.043 | 0.009 | 0.006 | 0.004 |

The effects of *bx-7AS* (functional allele) on FHB resistance was further investigated using a double haploid (DH) population (198 lines) derived from Yangmai-16 (dysfunctional allele) and Zhongmai-895 (**Table 2**). FHB index was calculated in both field and glasshouse conditions. Under the field growing condition, three developmental stages (20 DPA, 24 DPA, 28 DPA) were analysed. Similar to the above association analyses, results on the DH population also revealed a significant association (*P* < 0.009) of the functional *bx-7AS* allele with a lower FHB index, which decreased by 23.15%, 21.32% and 19.35 % for 20 DPA, 24 DPA and 28 DPA, respectively. For the glasshouse growing condition at 21 DPA, significant association (P = 0.043) of the functional *bx-7AS* allele with a lower FHB index was also observed, although leading to a relatively milder decrease (12.57%) on the FHB index. Taken together, association analyses showed that the functional alleles of ALP genes *bx-7AS* were significantly associated with FHB resistance.

### Overexpression of *TaALP-bx-7AS* gene in transgenic wheat lines revealed decreases in FHB symptoms

In order to further assess the involvement of the *ALPs* gene in wheat resistance to FHB, we generated transgenic wheat plants that had *bx-7AS* gene overexpressed. Two *ALPs* overexpression (*ALPs*ox) lines in the wheat cv. Fielder background were produced. *ALPs*ox #1 and #2 lines were inoculated together with wheat cv. Fielder and found elevated resistance to FHB. As shown in the **Table 3**, from 7 to 14 DPA, the rate of infected spikelet number was significantly decreased in the *ALPsox* lines compared with the control, suggesting slower FHB symptoms development. Similar patterns were found from 14 to 21 DPA in the *ALPsox* Lines. These two *ALPs*ox lines FHB spreading are reduced when compared with the control. Thus, results of the overexpression experiments strongly suggest that the *bx-7AS* ALP gene functions as a disease resistant components to FHB resistance.

**Table 3.** Relative increasing rate of infected spikelet number in transgenic and control wheat

| Wheat Lines | Relative increasing rate of infected spikelet (%) |  |
| --- | --- | --- |
|  | 7~14 DPA | 14~21 DPA |
| Control | 62.02 a | 44.50 a |
| 7A1-1 | 21.11 bc | 36.61 ab |
| 7A1-2 | 49.49 ab | 28.06 abc |
| 7A2-2 | 16.67 c | 21.18 bc |
| 7A2-4 | 21.42 bc | 7.87 c |

## Discussion

### Promoter significance of *TaALP genes*

FHB-responsive JA signalling regulated gene expression is immediate and conform in the resistance of wheat cvs. (*20, 63*). Many cysteine-rich antimicrobial peptides (AMPs) were found to be up-regulated by JA signalling, and were reported to be synthesized in healthy plants to maintain normal plant development (*64*) as well as functioning as a primary protection against diseases and pests(*65*). Meanwhile, an increased ethylene production contributes to wheat FHB resistance (*18*). Indeed, indications for an active ET signalling were found in the FHB-attacked resistant wheat transcriptome (*20*). In addition to the presence of JA- and ET-mediated general antifungal defences, a second line of defence was found to be based on a FHB-responsive and targeted suppression of relevant *Fusarium* virulence factors, such as proteases and mycotoxins (*20*). *Fusarium* subtilisin-like and trypsin-like proteases are released in infected wheat kernels mainly to disrupt host cell membranes during necrotrophic intracellular nutrition, both the plant protease inhibitor proteins and subtilisin-like and trypsin-like proteases of *F. graminearum* and *F. culmorum* have already been proven in the cereal grains (*20, 66*).

Previous studies showed that downstream PR genes are usually regulated by different signalling hormones, *ChiI* and *GluD* was located downstream of the ET pathway, *PR.10* was allocated downstream of the JA pathway, and *PR.1* was allocated downstream of the SA pathway (*67, 68*). Moreover, *in vitro* antifungal assays confirmed that the purified wheat *ChiI* and *GluD* proteins could inhibit the hyphal growth of *F. graminearum* (*69*). Base on promoter analysis, promoter regions of *TaALP* genes, were pathogen inducible, can be induced by disease related transcriptional factors (TFs), such as *TGAC-bZIP*/*ATB-bZIP* (Figure 1A). For example, at *ALP* gene loci on chromosome 7A, 31 and 4 motifs were identified for ATB_bZIP and TGA1_bZIP, respectively (Figure 1A). Further, promoter region motif annotation illustrated that *TaALP* genes have motifs related to SA and JA, ET signalling regulation (**Figure 1A**). A few ET/JA related motifs were compounded by pathogen responsive motifs, indicating that *TaALP* genes are likely to be regulated by ET/JA and located downstream of ET/JA signal pathway (**Figure 1A**). For example, at *ALP* gene loci on chromosome 7A, 137 pathogen related motifs were identified, 75 and 100 motifs were identified for ET and JA, respectively, yet with only 23 motifs identified for SA signal (**Figure 1A**). This results suggest that *TaALP* genes are likely to be induced by a JA or ET signal, and ET/JA signal pathway might act in a synergistic or opposite manner with SA signal pathway to confer FHB resistance. Transcript profile analysis by q-RT-PCR indicated that the transcripts of *TaALP* genes in wheat are induced by pathogen infection. Further, we have seen that the expression pattern of *TaALP* genes is similar to some PR genes (*Taxi III* and *PR.1*) under *F. graminearum* infection conditions. As shown in RT-PCR analysis above, the expression of *TaALP* was induced rapidly and dramatically by exogenous *F. graminearum* at 7 DAP for wheat cv. Spitfire and DH line 241, declined later on, whereas the expression of *TaALP* genes was dramatically induced at 13 DPA and the maturity stage for wheat cv. Mace. The transcriptional differences might be a direct results of differences in the ET/JA signalling. The dual peak, indicated by 13 DPA and maturity upregulation, might be caused by regulation of the expression of *TaALP* genes by other TFs regulated by ET/JA, or interactively regulated by other hormones, such as SA. Cross-talking between different signalling pathways might either activate or suppress the PR genes transcription (*70-72*). Taking these results together, *TaALP* is potentially involved in wheat defense response to *F. graminearum* through the ET/JA pathways.

In summary, the results of q-RT-PCR analyses showed that under *F. graminearum* infection, wheat grain dramatically increased the transcript levels of *TaALP* genes. And in addition, in the protein level, *in vitro* antifungal assays of recombinant protein products of *TaALP* genes gave evidence to their toxicity against hyphae growth of *F. graminearum* (**Figure 5D, E**). Most importantly, our results demonstrate that ALPs are directly involved in resistance to *F. graminearum* in wheat. ALPs, as a so far unknown family of antifungal proteins, can be used to breed wheat lines with increased disease resistance. Other researches were done on transformation of bread wheat by the transfer of cDNAs encoding differently acting antifungal proteins (*67, 73-76*). According to Ferreira *et al*. (*64*), overexpression of defense protein genes in the living host cells form a zone surrounding the most advancing hyphae as they allow a continuous supply with antifungals onto the intercellular hyphal tips. *TaALP* could be used as a candidate to improve crop resistance to *F. graminearum*. To our knowledge, this is the first time that ALPs, belonging to the seed prolamin superfamily with a gliadin domain, are reported to act as defense proteins against pathogens.

### Gliadin domain components display antifungal effects

ALPs contain either one or two gliadins domains (PF13016). Such a domain was also found to be characteristic of puroindolines, gamma and alpha gliadins and LMW glutenins. Similar to the pFam classification, ALPs has bifunctional inhibitor/plant lipid transfer protein/seed storage helical domain (*Bifun_inhib/LTP/seed_sf*), based on the InterPro classification. This represents a homologous superfamily of structural domains consisting of 4 helices with a folded leaf topology and forming a right-handed superhelix. Prolamin superfamily protein’s function may relate to protease inhibition or involvement in plant defence. As discussed in Juhasz, et al. 2018 (*77*), the hydrophobic-seed domain containing proteins include, cortical cell delineating (*78*), hydropho-seed domain containing protein (*79-83*), glycine-rich protein (*84-86*) and proline-rich protein (*87, 88*), which are found to be included in the plant defence system and have antifungal properties. Lipid transfer protein (*39, 40, 89-91*) and non-specific lipid transfer protein (*92*) have a LTP-2 domain, and have antifungal properties. Alpha-amylase/trypsin inhibitor (*93, 94*), Grain softness protein (*95*), Puroindoline (*96, 97*), Alpha gliadin (*98*) all contain a Tryp-alpha-amyl domain, and are known antifungal proteins. Meanwhile, Puroindoline, Alpha gliadin, LMW glutenin, Gamma gliadin, ALPs have a Gliadin domain, yet till now, the exact function is unclear. 19KDa Globulin (*99, 100*), Small cysteine-rich protein (*101-103*) belongs to the Domainless Cys-rich proteins, are involved in plant defence. While Omega gliadin and HMW-GS are Domainless Cys-poor proteins, were not reported to have disease resistance properties. Our study is the first-time proteins with gliadin domains that are also characteristic in gamma and alpha gliadins and LMW glutenins are described with a defense related function and this highlights the possible involvement of the gliadin domain in plant immunity and biotic stress mechanisms.

### Physio-chemical properties of ALPs protein identified in wheat cvs. Mace and Spitfire

Procedures for sequentially extracting and recovering protein fractions from small flour samples were described as reported previously (*48*). The NaI-propanol solution solubilized almost all the gliadins, albumins, and globulins, along with traces of glutenin (*48*). The present investigation has identified water and salt soluble proteins using three different approaches (RP-HPLC, SDS-PAGE, and MALFI-TOF) (**Figure S1, Figure 2, Figure S2A-F, Figure S3A-F, File S1, Table S2**). Analysis of the protein fractions by a combination of RP-HPLC followed by SDS-PAGE analysis along with protein reference maps developed by use of protein peptide sequencing or mass spectrometry, makes it possible to separate and identify most of the abundant proteins. These proteins include alpha-amylase and protease inhibitors (*104*), high molecular weight albumins (*105*) and other non-storage groups and enzymes which have specific synthetic, metabolic, regulatory, or protective roles in the plant (*106, 107*). Apart from this, some high molecular weight albumins and certain globulins (triticins) are considered to have a storage function (*108*).

RP-HPLC followed by SDS-PAGE of the albumin/globulin fraction demonstrated that it was highly enriched in ALPs. Identification of the ALPs was done by molecular mass based on MALDI-TOF analysis. In our analysis, when taken into all the obtained fragmentation patterns and aligned with the respective ALP amino acids sequences, most of the bands can be resolved. On the contrary, identification of *α-/β-* and *γ*-gliadins and LMW-GS by mass spectrometry tends to give low expectation score, due to the repetitive motifs in the N-terminal regions and proline-rich pattern, which are hard to digest with trypsin (*48*). In the case of ALPs identification, the only problem is the accurate determination of the homeologous proteins from 7A, 4A, and 7D, due to their highly similar amino acid sequences (> 93%). We did not attempt to resolve completely the individual subunits of this highly complex mixture. Likely, the many individual proteins in the region with apparent molecular masses from 33000 to 48000 Da (mainly gliadins and LMW-GSs) were not resolved by SDS-PAGE, may be due to overlapping of fractions because baseline separations were not achieved by RP-HPLC (**Figure S1**). Some of the individual ALPs are clearly resolved at apparent molecular masses of 17000 to 32000 Da, and consist of chromosomes 7A/4A/7D (**Figure 2, File S1, Table S2**). Protein bands below 16000 Da included LMW-albumins, such as members of the complex *α*-amylase inhibitor and *α*-amylase-trypsin inhibitor families that range in mass from 13000 to18000 Da. Protein bands of the molecular mass range of 30000 to 32000 Da include the homologous chromosome 7A/7D/4A-encoded type b ALPs. Protein bands in this size range also include the *α-/β*- and *γ*-gliadins, grain softness proteins, and the LMW-GS. It is unclear whether the homeologous chromosome 4A-encoded by-4AL and C-terminal by-4AL resolved in the same bands as by-7DS and C-terminal by-7DS. A distinct band of chromosome 7D-encoded ay-7DS and chromosome 4A-encoded ay-4AL, both with a molecular mass of approximately 18000 Da, were identified, respectively. The protein identification indicated that our method gave considerable overlap of protein types. For example, the ay-7AS proteins were eluted in peaks 5-7. The ay-4AL proteins were eluted in peaks 8-10 of wheat cv. Mace. The type b ALPs (bx-4AL/7DS) were detected in peaks 8, 9, 10, 11, 12, 13, 15, 17, 20, 21 of wheat cv. Mace. This was typical of most fractions in our study, which consisted of analysis of overlapping fractions corresponding to almost the entire area of the chromatogram (ALPs region). The different ALPs subunits have variant physio-chemical properties, ALPs ay, by, and bx subunits are similar to protease inhibitors like *α*-, *β*- amylase/subtilisin-inhibitors and serpins, triticins, while ALPs ax subunits are more similar physio-chemical properties as avenin-3, gliadins and LMW-GSs.

The identities of individual proteins separated by RP-HPLC here were also correlated with those of proteins resolved by others work. Shewry et al. (*109*) characterized certain seed albumins from different wheat species by N-terminal sequencing and found that several belonged to the trypsin/alpha-amylase inhibitor family. By using wheat null genetic lines, Singh and Skerritt (*110*) has established the location of several of their genes on individual chromosomes for albumin and globulin proteins. SDS-PAGE analysis of water-soluble proteins indicated the chromosomal location of polypeptides and proteins of different molecular weight were assigned on and 1D, 2A, 2B, 2D, 3AL, 3BS, 3DS, 4AL, 4BS, 4DS, 4DL, 5DL, 6DS, 7BS or 7DL (*110*). In our study, besides the identification of ALPs on 7DS, 4AL, and 7AS, it is also displayed in our analysis that other water- and salt-soluble proteins were located to chromosomes 1A/1B/1D (Avenin-3, Gamma-gliadin B, *γ-*gliadins and LMW-GS), 2A/2B/2D (alpha-amylase/subtilisin inhibitor), 3A/3B/3D (Alpha-amylase inhibitor), 4A/4B/4D (Alpha-amylase/trypsin inhibitor CM3), 5A/5B/5D (Grain softness protein), 6A/6B/6D (*α-/β-* gliadins), 7A/7B/7D (60S acidic ribosomal protein, Alpha-amylase/trypsin inhibitor CM2). Immunological and N-terminal sequencing characterisation identified most of the water-soluble proteins belonged to a family of *alpha* -amylase inhibitors, serine carboxypeptidase III homologous protein (*106*). Salt-soluble proteins matched with barley embryo globulins, other proteins include, lipid transfer protein (LTP), peroxidase BP-1 precursor and histone H4 proteins (*106*). The protein sequences could also potentially be used for making antibody or DNA probes for use in selection in breeding programmes. Information on the genetics and regulation of this fraction of proteins is necessary to understand their role and function in the grain. It is likely that proteins with similar physio-chemical properties are accumulated in the same fraction. The ALPs identified together with other antifungal proteins in albumin and globulin fraction might indicate similar antifungal functions.

### Temporal and spatial expression of *TaALP* gene under fungal infection

Plants induce defense responses against pathogen invasion which include activation of the SA-, JA-, ET-mediated defense pathways, which in turn increase reactive oxygen species (ROS) production, phytoalexin accumulation, Hypersensitive Response (HR), and/or upregulation of pathogenesis-related (PR) protein expression (*111*). Phyto-oxylipins comprising antimicrobial peptides and defence-signalling molecules such as JA, together with cysteine-rich PR genes indicate an induced antifungal defence mechanism (*20*). There is increasing evidence that members of the prolamin superfamily may play important roles in responding to biotic and abiotic stresses (*112-114*).

To understand the defense mechanisms of wheat grain ALPs, it is necessary to identify wheat *TaALP* genes and study their functions in the defense response to pathogens. In this study, we isolated and characterized pathogen induced *TaALP* genes in wheat, *TaALP*, whose transcript peak showed more rapid and stronger response to challenge with *F. graminearum* in the wheat cv. Spitfire and DH line 241 than in that of the wheat cv. Mace (**Table 1**). Our observation showed that disease symptoms in wheat cv. Mace were more severe than wheat cv. Spitfire and DH line 241 under *F. graminearum* infection. Meanwhile, *TaALP* genes induction, as well as the two *PR* genes (*Taxi III* and *PR.1*) showed earlier induction in Spitfire wheat and DH line 241 than in wheat cv. Mace. However, even the lower induction ratio at 7 DAP for wheat cv. Mace was considered relevant due to the strictly suppressed expression in the susceptible genotype. The significant >80-fold inductions at maturity stages for Mace wheat in the infected spike tissues was observed and may be an indication of delayed hormone regulation of the susceptible wheat cv. No gene expression was verifiable in spike samples of wheat cv. Spitfire and Line 241 for some *TaALP* genes at maturity stages. In the first instance, the relative induction peak at 7 DAP for wheat cv. Spitfire and Line 241 are an indication of earlier response of fungal infection and was consistent with less infected spikelets observations for wheat cv. Spitfire and Line 241 (20-30% infected spikelets) than wheat cv. Mace (50% infected spikelets). Given that *TaALP* with distinct transcript kinetics following pathogen challenge play unique roles in the defense response, it is necessary to identify wheat *TaALP* genes with stronger and faster induction by the pathogen. Wheat cvs. Spitfire and Mace may have developed a strategy to increase induction of *TaALP* expression, as well as other PR genes to counter infection by *F. graminearum*.

*TaALP* genes were specifically highly expressed in developing wheat caryopsis with a peak around 10-18 DAP. And in our study, we found enriched transcripts in fungal infected grains. *TaALP* genes transcripts can be localized in transfer cells as well as aleurone cells in the infected caryopses (**Figure 4**). It is likely that expression of *TaALP* are greatly induced in the wheat endosperm, sub aleurone cells and embryos, transfer cells, as well as the pericarps (**Figure 4**), which confirms that it acts directly on reduction of pathogen spread, and most likely has a role in plant immunity. The transcripts, however, were not restricted to the basal transfer cells; they were transcribed in the upper halves of immature kernels like aleurone cells, as well as the seed endosperm itself, and embryo, as was evidenced by mRNA *in situ* hybridization of *TaALP* genes (**Figure 4**). These up-regulated transcripts are most likely to represent defences, such as trigger mechanisms or direct antimicrobial activities (*74*).

Developing seeds are strong sinks for nutrients produced in the maternal plant (*115*). In wheat and barley, transfer cells are a vascular bundle running along the length of the grain, through modified maternal cells in the nucellar projection, to the endosperm cavity that extends along the seed, in parallel to the vascular bundle (*116*). The expression of *TaALP* in the transfer cells as well as aleurone cells under pathogen infection is consistent with the evidence that endosperm transfer cells maintain a delicate balance between nutrients transportation and the need to impede the ingress of pathogens into the developing seed (*117*). Transfer cells are involved in delivery of nutrients between generations and in the development of reproductive organs and also facilitate the exchange of nutrients that characterize symbiotic associations (*116*). Since transfer cells play important roles in plant development and productivity, the latter being relevant to crop yield, understanding the molecular and cellular events leading to wall ingrowth deposition holds exciting promise to develop new strategies to improve plant performance (*116*).

Enriched *TaALP* genes transcripts can also be localized in developing embryo around 15-18 DAP, which also indicate another category that might be of great interests. Proteomic study have indicated that proteins induced in response to infection are proteins involved in protein synthesis, folding and stabilization, as well as proteins involved in oxidative stress tolerance, and PR proteins in tissues of the fungal-infected germinating embryo (*118*). Lectin are known to have antifungal properties and are actively involved in plant defense, are expressed at low levels in the developing embryo together with the more abundant seed storage proteins (*119*).

### *In vitro* antifungal function of ALPs and allelic effect of ALPs on field FHB index

Further heterologous bacterial expression confirmed that TaALP could significantly reduce fungal hyphae growth *in vitro* (**Figure 5D**). As was illustrated in our *in vitro* antifungal activity test against *F. graminearum* fungal growth, EC_50_ values suggested that different paralogs of ALPs might differ in their toxicity (**Figure 5E**). In the transcriptional study under *F. graminearum* infection, *TaALP* encoded proteins belonging to the type a subgroup (ax-7AS/7DS and ay-7AS/7DS) can be induced at earlier stages, while most of the type b subgroup (bx-7AS/7DS and by-7AS/7DS) of the *TaALP* family can be induced at late grain filling stages under pathogen infection. These findings support the hypothesis that ALPs, might reduce certain protease activity of virulent pathogens as shown in the inhibited pathogen growth and spreading with much earlier induction in wheat cv. Spitfire and DH line 241. Therefore, as new members of the PR families and one of the many antifungal components, ALPs, are likely to play an important role in SAR and defense responses to *F. graminearum* infection initially, and some ALPs members induced at late grain filling stage could protect seed against this pathogen during germination.

*TaALP* genes encode prolamin superfamily member proteins that bear both antifungal properties while still maintaining the potential nutrient reservoir activity underpinned by typical storage proteins. Both in qualitative and quantitative aspect, ALPs might be of minor contribution to the total nutrient reservoir activity compared with glutenins, gliadins, and some HMW albumins and globulins (*120*). The induction of *TaALP* under pathogen infection, illustrated that they could act as pathogen resistant proteins that combat pathogen attack and assist plant survival under biotic stress. In our study, we hypothesized that ALPs are composed of a few gene members that work synergistically during grain filling, which is similar to the peptidase inhibitors of the defensin family (PR-12), which make up the third class of continual up-regulated AMPs (*121*). In particular, these constitutively expressed genes are supposed to contribute to non-host resistance (*122*). As was evident that most of the up-regulated cysteine-rich AMPs in resistant wheat cvs. have shown expression values that were independent of the treatment (*121*), but were lower or absent in the susceptible wheat cvs., which helps explains the differential transcripts indicated by the FHB induced expression of *TaALP* genes in wheat cv. Mace and Spitfire (**Table 1**). It is likely that ALPs, together with other AMPs, act synergistically in a generalized non-specific defence providing a basal protection (*20*). AMPs transcribed at a constant level are known key components of an immediate defence against invading pathogens, and many proteins are pathogen-inducible, for example, in leaves were found to be constitutively present in storage tissues, such as seed (*20*). This explains why wheat cv. Spitfire and the DH Line 241, which displayed high level expression of *TaALP* at 7 DAP, can suppress infection much more quickly than wheat cv. Mace, which were induced later (**Table 1**). Liu and others (*73*) described that genetically modified plants overexpressing certain antifungal peptides, would provide a promising alternative to improve overall resistance to *Fusarium* pathogen in wheat. Moreover, over-expressions of pathogen-inducible promoters directly targeting the infection sites or the most vulnerable tissues provides an approach to reducing the pathogenesis of the biotrophic *F. graminearum* fungi in colonized tissues (*123*). These possibilities illustrate that the promoters of *TaALP* genes can be of research interest and that ALPs can be used as novel antifungal peptides.

Variation in the amino acid sequences of the b-type proteins between the species suggest that they could provide a source of variation for wheat improvement (*124*). Whether all the *TaALP* genes function individually or collectively in conferring the observed broad-spectrum resistance is unknown. Nevertheless, *TaALP* are discussed as candidates for an improved resistance strategy against grain-infecting fungal pathogens and our results from RT-PCR analyses do not contradict these considerations. Large scale *Fusarium* phenotyping (FHB index) indicated that resistance was associated with allelic variation (bx-7AS allele) (**Table 2**). We propose that *TaALP* genes and their alleles are important in *Fusarium* resistance and can be utilized in breeding programs. We think that breeding for the presence of highly expressed *TaALP* genes can increase *Fusarium* resistance. Ov

### ALPs inhibition hypothesis

ALPs has peptides possibly involved in myristylation, phosphorylation, or glycosylation, or act as ligands of IG-MHC (Immunoglobulin major histocompatibility complex) (**Figure 1B**).

The synthesis of antimicrobial proteins is not restricted to plant species but seems to be ubiquitous in nature (*125*). The mould *Aspergillus giganteus* secretes the antifungal protein *Ag*-AFP, which displays inhibitory effects on the growth of phytopathogenic fungi (*125*). It is suggested that toxicity comes from an interaction of positively charged sites of the small protein with negatively charged phospholipids of susceptible fungal membranes (*126*). The outcomes of previous research indicated that pectin mythel esterase (*PME*) genes code for enzymes that are involved in structural modifications of the plant cell wall during plant growth and development (*127*). Some *F. graminearum* extracellular proteins, including pectin-degrading oligogalacturonases, can act as elicitors of defence reactions (*14*). A transgenic wheat line carrying a combination of a wheat *β*-1,3-glucanase and chitinase genes enhanced resistance against *F. graminearum* (*128*). Recently published approaches such as, expression of a pectin methyl-esterase inhibiting proteins (*129*) and polygalacturonase inhibiting proteins (PGIPs) (*34*), an antifungal radish defensin (*130*), a truncated form of yeast ribosomal protein L3 (*131*) and a phytoalexin Zealexin (*132*) have all shown to provide quantitative resistance against FHB. Defensins are a class of PR proteins with structurally related small, highly basic, and cysteine-rich peptides, which display broad-spectrum in vitro antifungal activities (*133, 134*). Maldonado et al. (*89*) demonstrated that LTPs, members of the prolamin superfamily, could either be a co-signal or act as a translocator for release of the mobile signal into the vascular system and/or chaperon the signal through the plant. Increased studies suggest that LTPs may be active defense proteins as biological receptors of elicitins, and play a significant role in activation of SAR mediated signalling pathways (*90, 135, 136*). Wheat contains three different classes of proteinaceous xylanase inhibitors (XIs), i.e. *Triticum aestivum* xylanase inhibitors (TAXIs) xylanase-inhibiting proteins (XIPs), and thaumatin-like xylanase inhibitors (TLXIs) which are believed to act as a defensive barrier against phytopathogenic attack (*33*). The up-regulation of thaumatin-like protein (TLP) is also observed, which can inhibit hyphal growth and/or spore germination of various pathogenic fungi through a membrane permeability mechanism or through degradation of fungal cell walls by *β*-1, 3 glucan binding and endo-*β*-1,3-glucanase activity (*32*).

Plant seeds, including cereal grains, contain numerous small protein inhibitors of proteinases (*104*). Some are efficient inhibitors of subtilisin-/chymotrypsin-like proteinases from microbes of insects, and it is more convincing now that they participate in an integrated broad spectrum defense system against invading fungal or insect pests (*137*). In the yeast two hybrid study, we found that ALPs are mostly like to interact with metacaspase (TaMCA4), which is a cysteine proteinase (**Figure 6**). In wheat and barley, homologous cysteine proteinases with optimal activity slightly below pH 5 play a central role in degradation of the prolamin storage proteins during germination (*138*). A highly possible hypothesis is that, ALPs, with the special cysteine rich structure of gliadin domains, are similar to the function of certain alpha-amylase inhibitors or serpins, and are likely to be toxic to fungal membranes. Amylase inhibitors/serpins act as suicide substrate inhibitors against certain proteinases, and the reactive centres of major serpins resemble the glutamine-rich repetitive sequences in prolamin storage proteins (*α*-, *γ*-, and ω-gliadins and the LMW and HMW subunits of polymeric glutenin) of wheat grain (*139, 140*). ALPs, like the well-known serpins, as baits, are likely to attract the amylase/trypsin/serine protease/cysteine aspartic protease by the glutamine-rich loops (mainly polyQs) between any of the four alpha-helices, and by inhibiting the peptide hydrolysis process, that of the protease function can be inhibited. Meanwhile. ALPs are also likely to have a weak interaction with N-myristoyltransferase (TaNMT) (**Figure 6**), this results indicate that the myristolation events for ALPs are highly possible, which lead to the PTM of certain ALPs. And most importantly, ALPs are able to interact with beta-glucosidase of *F. graminearum*. As is known, pathogen beta-glucosidase are able to hydrolyse cell wall components of host plants, the antifungal function of ALPs might suggest that they are able to inhibit the beta-glucosidase activity.

The storage tissues of plant seeds are attractive host for many pathogens. Evolutionary adaptation of the proteolytic system of some pathogen to efficient degradation of the abundant glutamine- and proline-rich repetitive structures of the cereal grain prolamins seems likely to have occurred. Here we have shown that the reactive centres of wheat grain ALPs contain unique glutamine-rich sequences resembling repetitive sequences of other wheat prolamins. A working hypothesis for further studies to elucidate the functions of the grain ALPs might be that the reactive centre loop sequences have evolved into a complement of baits for irreversible inactivation of cysteine proteinases, *etc*. from infection fungal, resulting in reduction of damage to seeds and thus in their increased survival.

### Conclusions

For the first time, we report that a prolamin superfamily member gene that encodes a protein with gliadin domains is involved in defense against *F. graminearum*. In silico analyses indicated the presence of critical peptides in TaALPs that are active in the plant immune system. The promoter motif contains abundant PR responsive motifs and hormone motifs. Expression levels of *TaALP* genes were significantly up-regulated when induced by infection of the fungus *F. graminearum*. And bacterially expressed ALPs displayed significant antifungal activity against wheat fungus *F. graminearum in vitro*. Genome wide association study indicated that there were significant allelic effects of *TaALP* genes on FHB indexes. For the first time, we have performed an *in situ* hybridization of *TaALP* genes in the developing caryopses, and we found enriched transcripts in the transfer cells, aleurone, sub-aleurone cells, and embryo of wheat caryopsis with significant FHB symptoms. In conclusion, we propose that these *TaALP* genes fulfil a PR protein function and are involved in SAR.

---

**Supporting Information**

**Author information**

Corresponding Author

*E-mail: Tel: Fax:

The authors declare no competing financial interest.

**Acknowledgments**

**Figure S1.**
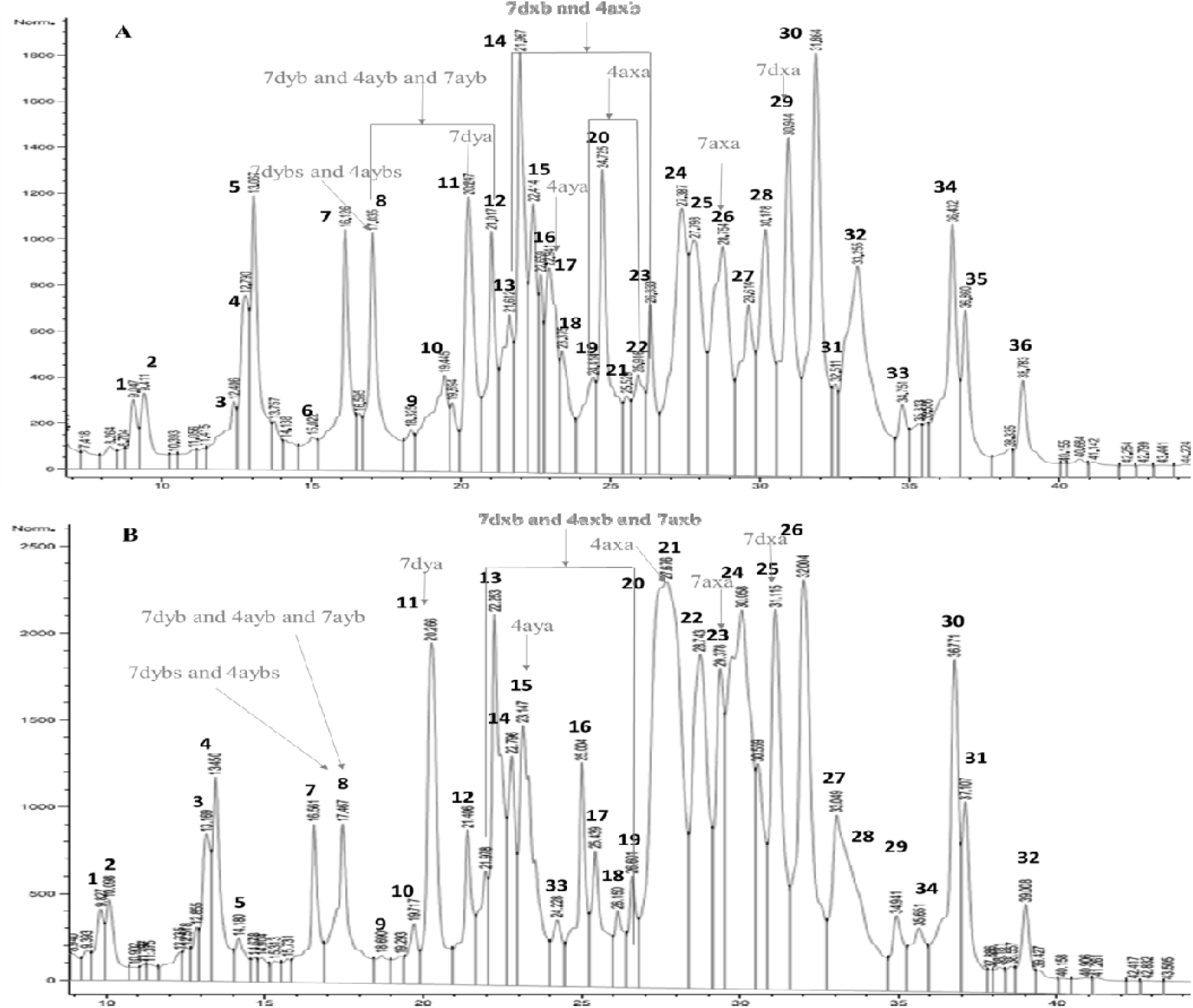
RP-HPLC analyses of albumin and globulin proteins in wheat. A: Mace; B: Spitfire.

**Figure S2A-F.**
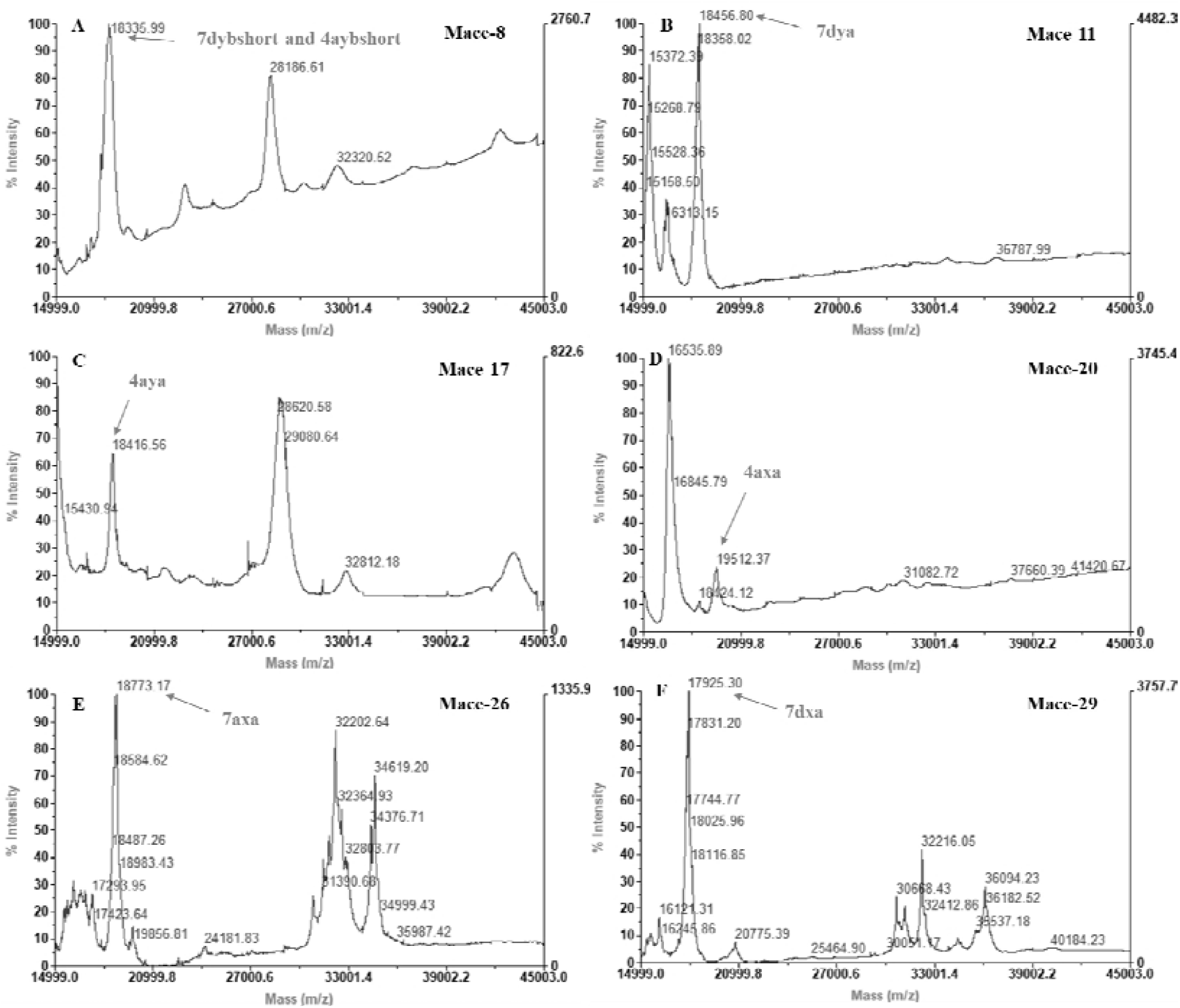
Maldi-tof analyses of ALP proteins present in wheat grain of Mace.

**Figure S3A-F.**
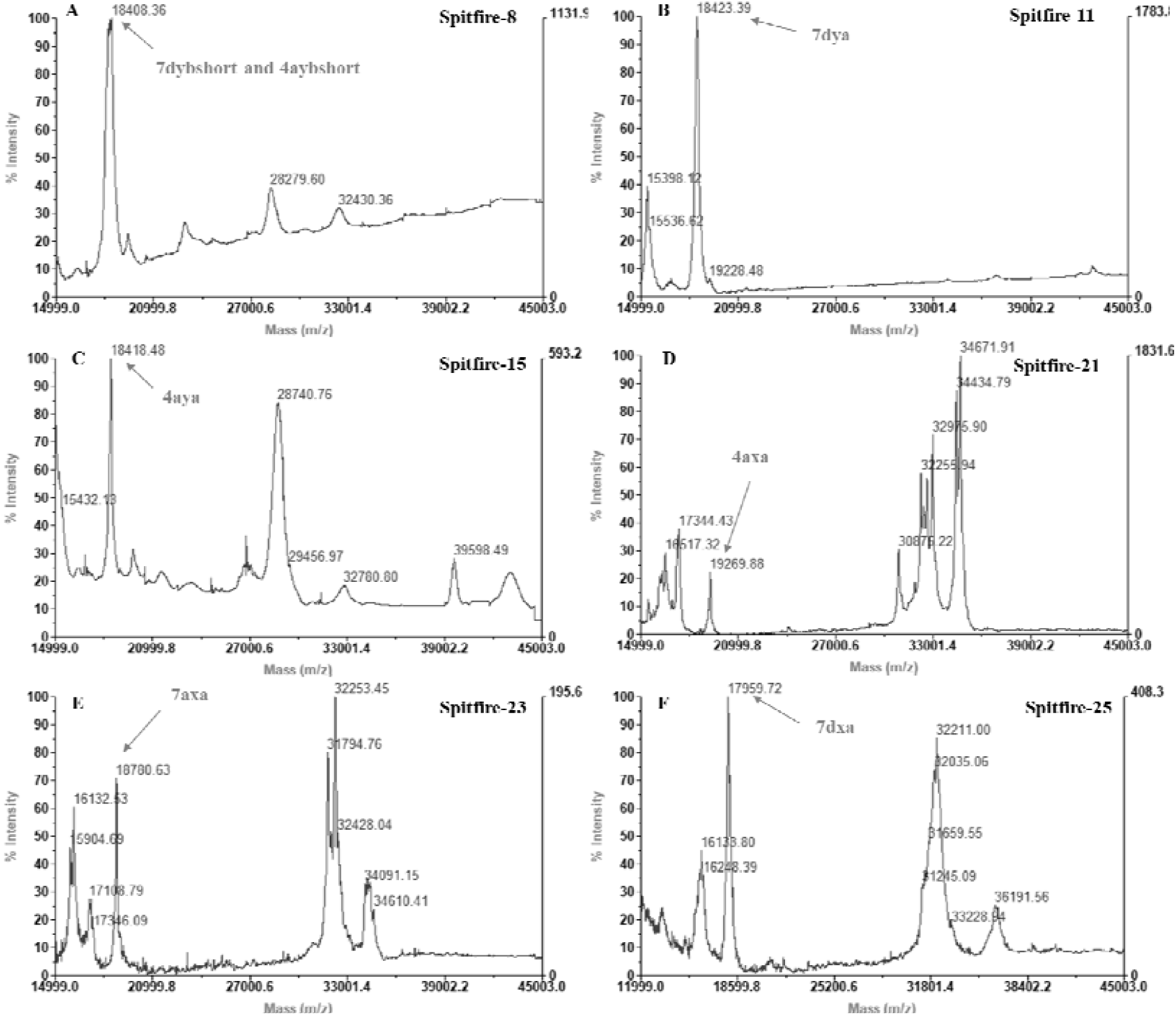
Maldi-tof analyses of ALP proteins present in wheat grain of Spitfire.

**Table S1.** The list of the retrieved motif related to pathogenesis.

|  | **Stress** | **Hormone** | **TF family** |
| --- | --- | --- | --- |
| **CANNTG** | pathogen | JA | bHLH MYC |
| **TGACG** | pathogen | SA | TGA bZIP |
| **TTGAC** | pathogen | SA | WRKY |
| **AAAGATA** | pathogen |  | GATA zinc finger |
| **AACGTG** | pathogen | JA | MYC |
| **ACGT** | Light, elicitor, pathogen | ethylene | bZIP |
| **ATCAT** | pathogen | JA | ATB bZIP |
| **AGCCGCC** | pathogen | Ethylene, JA | AP2/ERF |
| **CTCTT** | pathogen |  |  |
| **GCCGCC** | pathogen | Ethylene, JA | AP2/ERF |
| **GTAC** | Biotic, abiotic<br>stress | ethylene | SBP |

Motifs were collected from PLACE and PlantCARE databases. Hormone and TF family specific information was retrieved from the annotation.

**Table S2.**
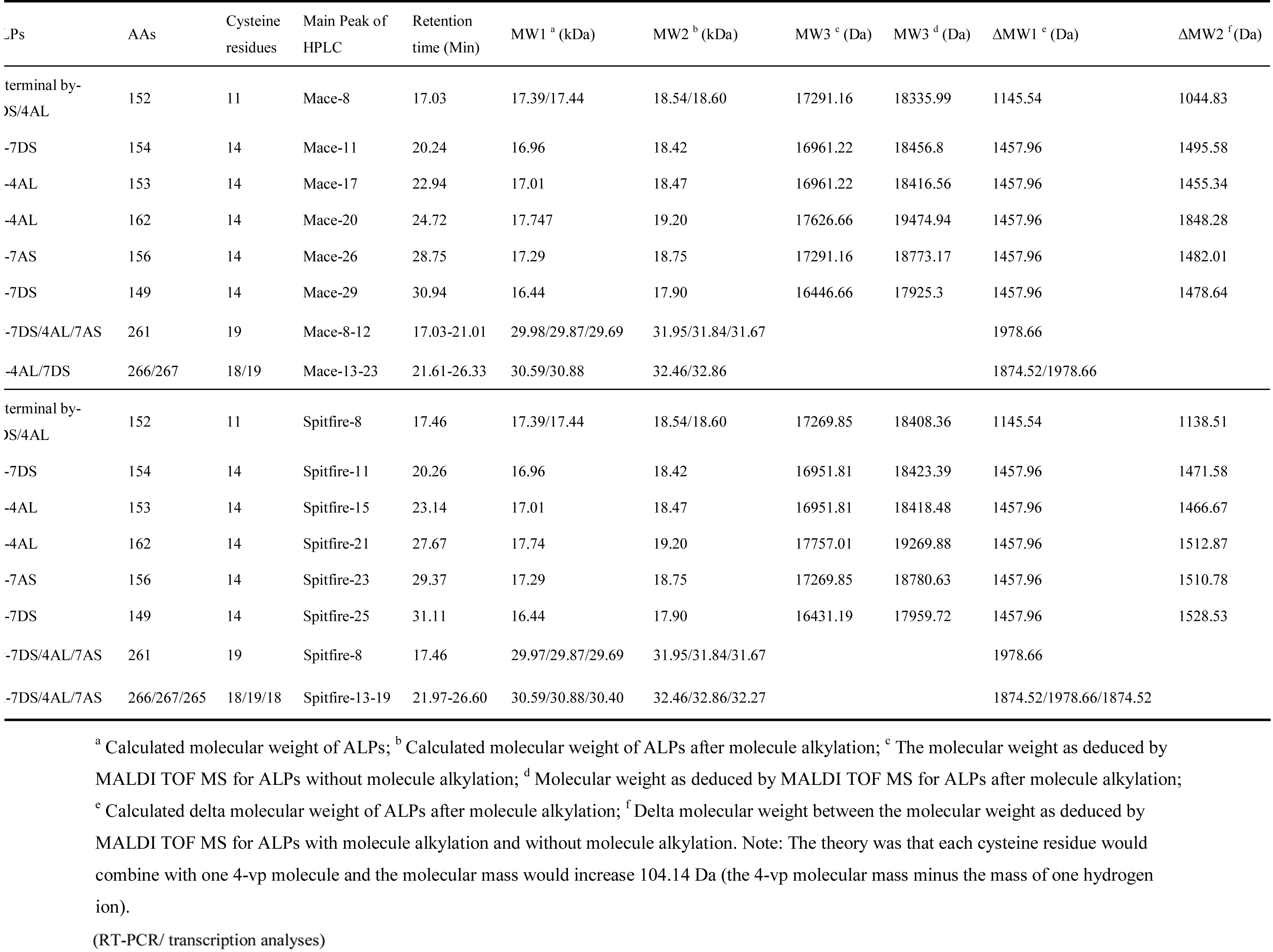
Sumary of the detected ALPs of with HPLC and MALDI-TOF.

**File S1.** Peptide sequencing results of wheat ALPs.

| SDS-PAGE bands | HPLC Peak | Retention Time | Sequencing Results | MW of MAILDI-TOF |
| --- | --- | --- | --- | --- |
| 1 | Mace-8 | 17.035 | C-terminal by-7DS/4AL | 18335.99 |
| 2 | Mace-8 | 17.035 | by-7DS/4AL/7AS | 28185.92 |
| 3 | Mace-9 | 18.325 | C-terminal by-7DS/4AL | 18337.51 |
| 4 | Mace-9 | 18.325 | by-7DS/4AL/7AS | 28185.92 |
| 5 | Mace-10 | 19.445 | C-terminal by-7DS/4AL | 18451.18 |
| 6 | Mace-10 | 19.445 | ay-7DS | 18451.18 |
| 7 | Mace-11 | 20.247 | ay-7DS | 18456.80 |
| 8 | Mace-11 | 20.247 | by-7DS/4AL/7AS |  |
| 9 | Mace-12 | 21.017 | by-7DS/4AL/7AS |  |
| 10 | Mace-13 | 21.612 | bx-7DS/4AL | 28736.77 |
| 11 | Mace-14 | 21.967 | bx-7DS/4AL | 28776.56 |
| 12 | Mace-15 | 22.414 | bx-7DS/4AL | 28617.72 |
| 13 | Mace-16 | 22.659 | bx-7DS/4AL | 28774.42 |
| 14 | Mace-17 | 22.941 | ay-4AL | 18416.56 |
| 15 | Mace-17 | 22.941 | bx-7DS/4AL | 28619.52 |
| 16 | Mace-18 | 23.375 | ay-4AL | 18428.36 |
| 17 | Mace-18 | 23.375 | bx-7DS/4AL | 28586.6 |
| 18 | Mace-19 | 24.414 | ax-4AL | 19515.57 |
| 19 | Mace-20 | 24.725 | ax-4AL | 19512.37 |
| 20 | Mace-20 | 24.725 | bx-7DS/4AL |  |
| 21 | Mace-21 | 25.529 | bx-7DS/4AL |  |
| 22 | Mace-24 | 27.387 | ax-7AS | 18845.22 |
| 23 | Mace-25 | 27.798 | ax-7AS | 18681.39 |
| 24 | Mace-26 | 28.754 | ax-7AS | 18773.17 |
| 25 | Mace-27 | 29.614 | ax-7AS | 18789.94 |
| 26 | Mace-28 | 30.178 | ax-7DS | 17990 |
| 27 | Mace-28 | 30.178 | ax-7AS | 18805.21 |
| 28 | Mace-29 | 30.914 | ax-7DS | 17925.30 |
| 29 | Mace-30 | 31.864 | ax-7DS | 18058.39 |
| 30 | Spitfire-8 | 17.467 | C-terminal by-7DS/4AL | 18408.36 |
| 31 | Spitfire-8 | 17.467 | by-7DS/4AL/7AS | 28236.77 |
| 32 | Spitfire-11 | 20.266 | ay-7DS | 18423.39 |
| 33 | Spitfire-12 | 21.406 | ay-7DS | 18447.6 |
| 34 | Spitfire-14 | 22.796 | ay-4AL | 18472.54 |
| 35 | Spitfire-14 | 22.796 | bx-7DS/4AL/7AS | 28759.24 |
| 36 | Spitfire-15 | 23.147 | ay-4AL | 18418.48 |
| 37 | Spitfire-15 | 23.147 | bx-7DS/4AL/7AS |  |
| 38 | Spitfire-15 | 23.147 | bx-7DS/4AL/7AS |  |
| 39 | Spitfire-16 | 25.004 | bx-7DS/4AL/7AS |  |
| 40 | Spitfire-18 | 26.15 | bx-7DS/4AL/7AS |  |
| 41 | Spitfire-18 | 26.15 | - |  |
| 42 | Spitfire-19 | 26.601 | bx-7DS/4AL/7AS |  |
| 43 | Spitfire-20 | 27.305 | ax-4AL | 18812.35 |
| 44 | Spitfire-21 | 27.676 | ax-7AS peptide 1 |  |
| 45 | Spitfire-21 | 27.676 | ?? |  |
| 46 | Spitfire-21 | 27.676 | ax-4AL | 19269.88 |
| 47 | Spitfire-22 | 28.743 | ax-7AS | 18787.16 |
| 48 | Spitfire-22 | 28.743 | ax-4AL | 19249.09 |
| 49 | Spitfire-23 | 29.378 | ax-7AS | 18780.63 |
| 50 | Spitfire-24 | 30.058 | ax-7AS peptide 2 |  |
| 51 | Spitfire-24 | 30.058 | ax-7DS |  |
| 52 | Spitfire-24 | 30.058 | ax-7AS | 18786.6 |
| 53 | Spitfire-25 | 31.115 | ax-7DS | 17959.72 |
| 54 | Spitfire-25 | 31.115 | ax-7DS | 18036.36 |
| 55 | Spitfire-26 | 32.004 | ax-7DS | 18188.99 |

